# Deep phenomics of African green monkey and Baboon PBMCs using a 30-marker spectral flow cytometry panel

**DOI:** 10.64898/2026.11.06.764971

**Authors:** Soumik Barman, Arsh Patel, Michael J. Buonopane, Ben Janoschek, Ravi Hingorani, David J. Dowling

## Abstract

Vaccines are the best way to prevent and combat diseases. Non-human primate (NHP) models have proven to be the gold standard model for studying the efficacy of certain vaccine candidates prior to human trials. Cell-mediated immunity serves as a surrogate marker for predicting vaccine efficacy. Despite the availability of deep-immunophenotyping panels for T cells in the peripheral blood of humans, their application to NHPs, particularly African Green Monkeys (AGMs) and baboons, remains limited. Notably, AGMs are valuable for HIV and COVID-19 research, while baboons are utilized for pertussis research as a natural infection model and vaccine trial. To overcome such limitation, using spectral flow cytometry, we developed a 30-parameter panel to profile major immune cell subsets (B cells, NK cells, DCs, monocytes) either in AGM or baboon PBMCs. We also profiled T cells, memory phenotypes, functional cytokines by tracking established Th1 markers IFNγ, TNF and IL-2; Th2 markers IL-4/5/13 and the Th17 marker, IL-17, activation-induced markers (AIMs) and FoxP3^+^ Tregs in both AGM and baboon PBMCs in steady state as well as in mitogen stimulated conditions. For AIMs, CD69, CD134 (OX40) and CD137 (4-1BB) are included and validated in both NHP species. This panel holds potential for exploratory analysis of the impact of these functional cytokines, AIMs, or Tregs on host immunity following infection or vaccination. The panel’s high level of customization empowers researchers to tailor it to their specific research needs.

## 1 Panel Presentation

### 1.1 Purpose and Appropriate Sample Types

Building a cross-species spectral flow cytometry panel is challenging because there’s a lack of data on the extensive validation of commercially available antibodies (Abs). OMIPs-52 (1) and 75 (2) serve as a comprehensive toolkit for profiling Th1/Th2/Th17 polarization in rhesus and cynomolgus macaque PBMCs. However, these cross-reactive Abs have never been validated in other NHP species. Hence, we validated a diverse range of clones among NHPs (AGM/baboons) in conjunction with humans (3) and developed a 28-color panel for the phenotypic analysis of antigen-specific T cells. This panel not only employs a conventional approach to profile Th polarization but also incorporates cytokine-independent AIMs. To dissect adaptive immune compartments, we phenotype Th1 cytokines (IFNγ, TNF, IL-2), Th2 cytokines (IL-4, IL-5, IL-13), and Th17 cytokine (IL-17) in effector CD4^+^ T cells. Validation of both Th polarization and AIM-specific markers (CD134, CD137, CD69) was conducted during ICS or nuclear staining. We profiled nuclear FoxP3^+^ Regulatory T cells (Treg) signatures in both AGMs and baboons. Notably, this is the first time that nuclear FoxP3^+^ Treg signatures have been profiled in NHPs in an OMIP. We developed this spectral flow panel to assess antigen specific T cells in AGMs triggered by a potential vaccine candidate to combat COVID-19 (ongoing research). We checked the cross-reactivity of this panel in pure olive or mixed olive-yellow baboons to further target innate/adaptive immune cell populations during the pertussis infection to understand the pathogenesis and disease clearance mechanisms (ongoing work). In parallel, we evaluate the cross-reactivity of this panel using human PBMCs and created its counterpart with a few fine-tuning adjustments (3). The human counterpart of this 28-color NHP panel served as a positive control during the panel development and extensive validation of Abs clones, stimulants, and reagents (3).

### 1.2 Background

OMIPs have been extended to encompass 45 colors (4) and beyond (5), surpassing the capabilities previously attainable solely through CyTOF. Despite such extraordinary achievement, phenotyping work using AGM and baboons was neglected. Currently, there is no optimized high-dimensional OMIP established to target functional cytokines (Th1, Th2, and Th17 cells) and Tregs in AGM or baboon models. Cytokine-independent AIM assays that identify antigen-specific T cells have never been properly validated using AGM or baboon PBMCs, despite these models being efficient for clinical trials. AGMs serve as valuable models for studying HIV (6,7) and SARS-CoV-2 (8–10). Baboons are utilized in pertussis research due to their natural susceptibility to the disease and their ability to mimic human symptoms (11,12), making them an ideal candidate for vaccine testing (13–15). Furthermore, the baboon model was also employed in COVID-19 research to evaluate the pathogenesis (16) and vaccine efficacy (17). This 30-parameter spectral flow cytometry panel was developed to assess the functional cytokine along with the AIMs signature by conventional T cells, which shape the host immune responses upon natural infection or vaccinations (18,19).

Using information from the database of published OMIPs (20) and prediction tools provided by the Nonhuman Primate Reagent Resource (21), we initially constructed a theoretical panel. We validated the locked-down clones simultaneously using PBMCs from AGM, baboons and humans and optimized the panel further (See **Section 3: Panel Design and Optimization**). Panel optimization was performed using a Sony ID7000™ spectral flow cytometer. This instrument, previously described by our group in OMIP-111 (22), is equipped with five lasers (355 nm, 405 nm, 488 nm, 561 nm, and 637 nm) and 147 fluorescence detectors. This panel has undergone extensive validation using various mitogens with NHP PBMCs. The T cell Receptors (TCR) dependent profile will be published as follow-up studies utilizing PBMCs from immunized or challenged NHPs.

### 1.3 Results

Figure 1 shows a representative comparison of major innate and adaptive immune cell populations, TCR-independent AIMs, Th Polarization, and FoxP3^+^ Treg signatures in (A-G) AGM and (H-N) olive baboon PBMCs after nuclear staining. We mirrored the gating strategy from OMIP-XXX-A (3). Readers can find the justification for the gating strategy in “Section 1.3: Results” of OMIP-XXX-A (3). FIt-SNE representation using OMIQ platform portrayed the cellular topography in NHPs (Figure 1G & N) after concatenation. Figure 1G shows the predominant island of memory CD8^+^ T cells (in brown) in the live CD45^+^ cellular landscape in AGMs. Such distribution is absent in baboons (Figure 1N). Enrichment of CD8^+^ T cells in AGMs is common (23) because of high viremia (7), which made them an ideal host for HIV research (6). Of note, AGMs are natural hosts of SIVs (6,23). Less enrichment of early NK cells was detected in AGMs (Figure 1B), likely due to high viremia. TCRγδ T cells are also less frequent in AGMs compared with baboons (Figure 1A & H), despite inter-individual variability among animals (Figure 7.4). The specific flow antibodies and their optimized titrations for this NHP-OMIP are documented in Table 2 and 4.

**Figure 1:**
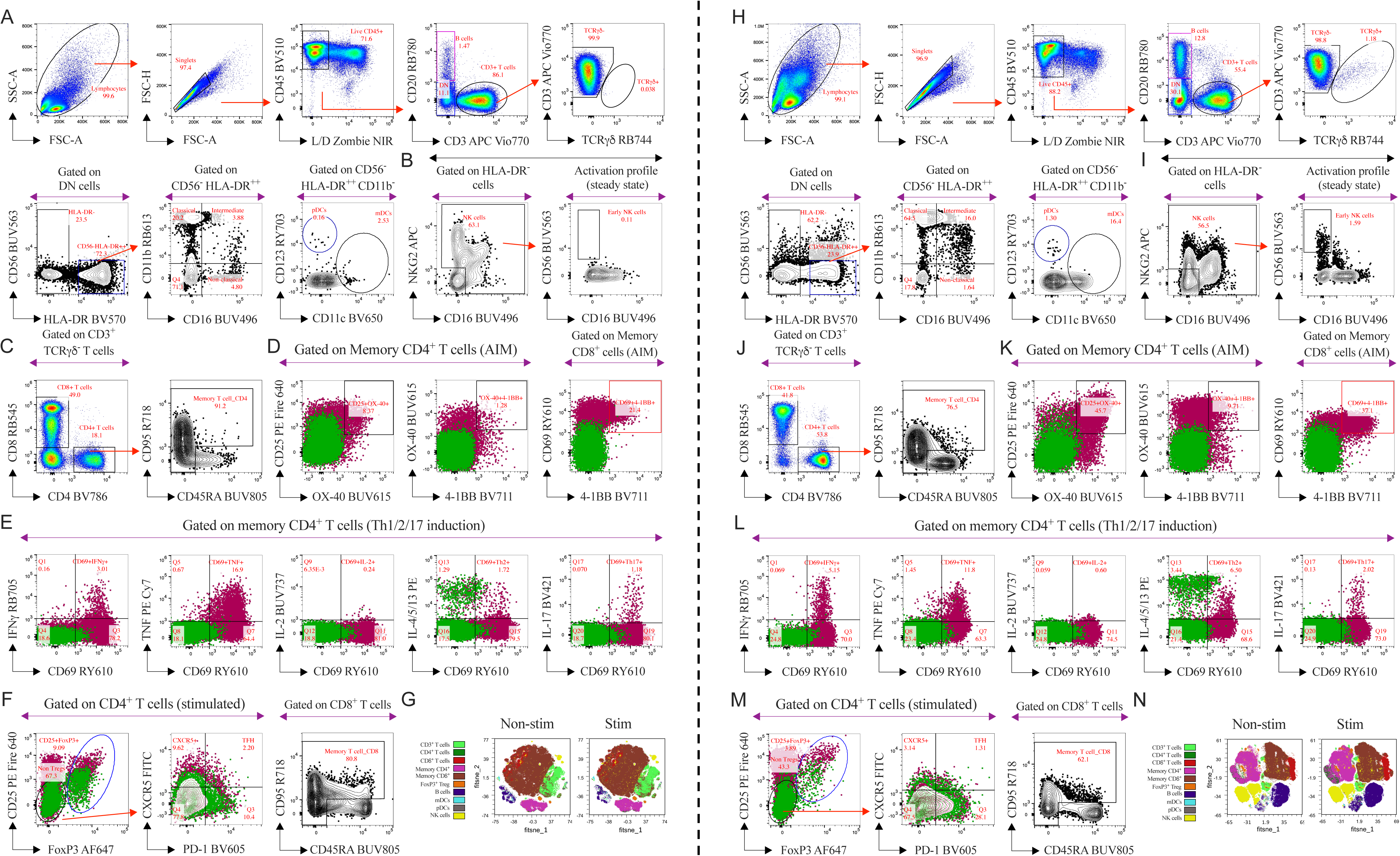
Representative gating strategy to profile activation induced markers (AIMs), Th1/2/17 polarization along with FoxP3^+^ Tregs and FIt-SNE overlays for the visualization of the immune cell populations using 30-parameter OMIP-XXX-B in cryopreserved PBMCs from (A-G) AGMs and (H-N) olive baboons. Sequential gating strategy was depicted by red arrows. (A & H) Following gating live CD45^+^ leukocytes, including doublet and dead cells exclusion, T and B cells were subsequently identified using CD3 and CD20, respectively. Non-B and T cells were further classified to monocytes, DCs and (B & I) NK cells. Conventional NHP CD3^+^ T cell compartment was dissected further to define TCRγδ T cells and non-TCRγδ cells were further phenotyped to portray (C & J) CD4^+^ and CD8^+^ T cells. Memory phenotype was assessed using CD95 and CD45RA expression. (D & K) NHP PBMCs were stimulated with Con A for 20 h and further phenotyped for the AIM profile in the memory T cell compartment according to the expression/combination of CD25, CD134 and CD137. (E & L) Intracellular Th1, Th2 and Th17 signatures were also captured upon Con A stimulation by memory CD4^+^ T cells. (F & M) In NHPs, CD4^+^ T cell subsets were also explored to phenotype CD25^+^ FoxP3^+^ Treg signatures or TFH (CXCR5^+^ PD-1^+^) signatures. (D-F & K-M) In representative bivariate plots, the AIM profile or Th1/2/17 polarization upon stimulation was portrayed by overlaying resting (in clover) subsets on its stimulated counterpart (in cayenne). Either 100K events of live CD45^+^ leukocytes from (G) each AGM (n=4) or (N) 120K events from each baboon (n=5) were used in concatenated visualization with FIt-SNE by the OMIQ platform. The FIt-SNE overlays captured the topography of the cellular landscape in NHP PBMCs in steady state or in Con A stimulated (20 h) conditions. For clarity, bivariate plots from the same NHP were presented either in steady state or in Con A stimulated conditions.

### 1.4 Similarity to Published OMIPs

Our OMIP is unique in its design, intended for dual species of NHPs: AGM and baboons. To date, only one OMIP (24) has been designed for AGM, but it did not dissect the T cell compartment, as its focus was primarily on B and plasma cells. Our OMIP is complex in nature, as it captures the TCR-independent AIM profile, Th polarization, and Treg subset simultaneously in both AGMs and baboons. OMIP-108 (25) has been designed for baboons PBMCs and is restricted to T cell compartment dissection and the AIM profile of CD69. We captured intracellular Th polarization and intranuclear Treg profile in baboons and expanded the AIM profile beyond CD69 based on CD134 and CD137. We adopted the clone NCAM16.2 for CD56 to phenotype NK cells and clone ICRF44 for CD11b as granulocyte and monocyte markers from OMIP-108 (25). We followed OMIP-5 (26) and 52’s (1) gating strategy to target functional cytokines on activated CD69^+^ T cells, but none of these OMIPs have a human counterpart. Notably, our OMIP has a corresponding human counterpart (OMIP-XXX-A, Table 1) (3).

**Table 1.** Summary table for the application of OMIP-XXX-B.

|  | **Summary Table** |
|---|---|
| Purpose | Deep phenomics of activation and functional cytokines of non-human primate T cells |
| Technology | Spectral Flow Cytometry |
| Application | Immunophenotyping of clinical and vaccine cohorts |
| Species | African green monkey (*Chlorocebus aethiops*), olive baboon (*Papio anubis*), and hybrid olive-yellow baboon (*Papio anubis/cynocephalus*) |
| Cell types | Cryopreserved PBMCs |
| Cross references | OMIP-5, OMIP-52, OMIP-108, OMIP-XXX-A<sup>3</sup> |

**Table 2.** Reagents for OMIP-XXX-B comprising the 30-parameter or the 28-color panel.

|  | **Specificity** | **Fluorochrome** | **Clone** | **Purpose** |
|---|---|---|---|---|
| 1 | CD3 | APC Vio770 | 10D12 | T cells |
| 2 | CD4 | BV786 | L200 | T cells |
| 3 | CD8 | Real Blue 545 | SK1 | T cells |
| 4 | CD11b | Real Blue 613 | ICRF44 | Monocytes, granulocytes |
| 5 | CD11c | BV650 | S-HCL-3 | Myeloid dendritic cells |
| 6 | CD16 | BUV496 | 3G8 | Monocytes, NK cells activation |
| 7 | CD20 | Real Blue 780 | 2H7 | B cells |
| 8 | CD25 | PE Fire 640 | M-A251 | Tregs |
| 9 | CD45 | BV510 | D058-1283 | Leukocytes |
| 10 | CD45RA | BUV805 | 5H9 | T cell memory |
| 11 | CD56 | BUV563 | NCAM16.2 | NK cells |
| 12 | CD69 | Real Yellow 610 | FN50 | Activation Induced Marker |
| 13 | CD95 | R718 | DX2 | T cell memory |
| 14 | CD123 | Real Yellow 703 | 7G3 | Plasmacytoid dendritic cells |
| 15 | CD134/OX-40 | BUV615 | L106 | Activation Induced Marker |
| 16 | CD137/4-1BB | BV711 | 4B4-1 | Activation Induced Marker |
| 17 | CD159a/NKG2a | APC | Z199 | NK cells |
| 18 | CD185/CXCR5 | FITC | MU5UBEE | TFH Marker |
| 19 | CD279/PD-1 | BV605 | EH12.1 | TFH Marker |
| 20 | FoxP3 | AF647 | 206D | Tregs |
| 21 | HLA-DR | BV570 | L243 | Dendritic cells, monocytes |
| 22 | IFNγ | Real Blue 705 | B27 | Th1 cells |
| 23 | IL-2 | BUV737 | MQ1-17H12 | Th1 cells |
| 24 | IL-4 | PE | 8D4-8 | Th2 cells |
| 25 | IL-5 | PE | JES1-39D10 | Th2 cells |
| 26 | IL-13 | PE | JES10-5A2 | Th2 cells |
| 27 | IL-17A | BV421 | N49-653 | Th17 cells |
| 28 | TCRγδ | Real Blue 744 | B1 | γδ T cells |
| 29 | TNF | PE Cy7 | MAb11 | Th1 cells |
| 30 | Viability | Zombie NIR | --- | Viability |

## 2 Technical Information

### 2.1 Instrument’s characteristics, setup and QC

This OMIP was developed using a Sony ID7000™ spectral cell analyzer equipped with five lasers (355 nm, 405 nm, 488 nm, 561 nm, and 637 nm) and 147 fluorescence detectors. The optical layout, fluorescence detector configuration, and fluorochrome distribution for the optimized panel are shown in Table 3. The instrumentation setup and daily QC procedures conducted by the Dana-Farber Cancer Institute (DFCI) cytometry core were documented in OMIP-XXX-A (3), which serves as a human counterpart to this panel. Intentionally, we bypass the documentation here to avoid exact repetition.

**Table 3.** The instrumental configuration for the Sony ID7000™ is presented, outlining the wavelength range of its spectral detectors. For each spectral detector across the five lasers, the first two columns list the channel band and its calculated center wavelength (median in nanometers). Detectors are named according to their position on the laser array, from the smallest to the largest wavelength. Notably, the UV and Violet lasers each utilize three distinct detectors with unique channel bands and center wavelengths, which are individually listed. Additionally, the panel illustrates the 28 fluorochromes, indicating the channel where each exhibits its peak fluorescence. Fluorochrome abbreviations are as follows: Brilliant UV (BUV), Brilliant Violet (BV), Real Blue (RB), Real Yellow (RY), Alexa Fluor (AF), and Red (R).

| **Detector Array** |  | **Ultraviolet** |  | **Violet** |  | **Blue** |  | **Yellow-Green** |  | **Red** |  |
|---|---|---|---|---|---|---|---|---|---|---|---|
|  |  | **355 nm** | **50 mW** | **405 nm** | **100 mW** | **488 nm** | **150 mW** | **561 nm** | **100 mW** | **637 nm** | **140 mW** |
| Channel Band (nm) | Center Wavelength (nm) | Detector | Fluorochrome | Detector | Fluorochrome | Detector | Fluorochrome | Detector | Fluorochrome | Detector | Fluorochrome |
| 360.5-390 | 375 | UV1 |  | - |  | - |  | - |  | - |  |
| 413.6-438 | **425** | - |  | **V1** | **BV421** | - |  | - |  | - |  |
| 418.5-446 | 430 | UV2 |  | - |  | - |  | - |  | - |  |
| 438-459 | 450 | - |  | V2 |  | - |  | - |  | - |  |
| 446-493.9 | **470** | **UV3** | **BUV496** | - |  | - |  | - |  | - |  |
| 449-480.2 | 465 | - |  | V3 |  | - |  | - |  | - |  |
| 493.9-506.1 | **500** | UV4 |  | V4 |  | **B4** | **FITC** | - |  | - |  |
| 506.1-518.3 | **510** | UV5 |  | **V5** | **BV510** | B5 |  | - |  | - |  |
| 518.3-530.3 | 525 | UV6 |  | V6 |  | B6 |  | - |  | - |  |
| 530.3-542.3 | 535 | UV7 |  | V7 |  | B7 |  | - |  | - |  |
| 542.3-554.2 | **550** | UV8 |  | V8 |  | **B8** | **RB545** | - |  | - |  |
| 554.2-566.1 | **560** | **UV9** | **BUV563** | V9 |  | B9 |  | - |  | - |  |
| 566.1-577.8 | **570** | UV10 |  | **V10** | **BV570** | B10 |  | YG10 |  | - |  |
| 577.8-589.5 | **585** | UV11 |  | V11 |  | B11 |  | **YG11** | **PE** | - |  |
| 589.5-601.1 | 595 | UV12 |  | V12 |  | B12 |  | YG12 |  | - |  |
| 601.1-612.6 | **605** | UV13 |  | **V13** | **BV605** | B13 |  | **YG13** | **RY610** | - |  |
| 612.6-624 | **620** | **UV14** | **BUV615** | V14 |  | **B14** | **RB613** | YG14 |  | - |  |
| 624-635.3 | 630 | UV15 |  | V15 |  | B15 |  | YG15 |  | - |  |
| 635.3-646.6 | **640** | UV16 |  | V16 |  | B16 |  | **YG16** | **PE Fire 640** | - |  |
| 646.6-657.7 | **650** | UV17 |  | **V17** | **BV650** | B17 |  | YG17 |  | R17 |  |
| 657.7-668.8 | **665** | UV18 |  | V18 |  | B18 |  | YG18 |  | **R18** | **APC** |
| 668.8-679.8 | **675** | UV19 |  | V19 |  | B19 |  | YG19 |  | **R19** | **AF647** |
| 679.8-690.8 | 685 | UV20 |  | V20 |  | B20 |  | YG20 |  | R20 |  |
| 690.8-701.6 | 695 | UV21 |  | V21 |  | B21 |  | YG21 |  | R21 |  |
| 701.6-712.4 | **705** | UV22 |  | **V22** | **BV711** | **B22** | **RB705** | **YG22** | **RY703** | R22 |  |
| 712.4-723 | **720** | UV23 |  | V23 |  | B23 |  | YG23 |  | **R23** | **R718** |
| 723-733.6 | 730 | UV24 |  | V24 |  | B24 |  | YG24 |  | R24 |  |
| 733.6-744.2 | **740** | **UV25** | **BUV737** | V25 |  | B25 |  | YG25 |  | R25 |  |
| 744.2-756.6 | **750** | UV26 |  | V26 |  | **B26** | **RB744** | YG26 |  | **R26** | **Zombie NIR** |
| 756.6-765 | 760 | UV27 |  | V27 |  | B27 |  | YG27 |  | R27 |  |
| 765-775.2 | **770** | UV28 |  | V28 |  | B28 |  | YG28 |  | **R28** | **APC Vio770** |
| 775.2-785.4 | **780** | UV29 |  | **V29** | **BV786** | **B29** | **RB780** | **YG29** | **PE Cy7** | R29 |  |
| 785.4-795.5 | 790 | UV30 |  | V30 |  | B30 |  | YG30 |  | R30 |  |
| 795.5-805.5 | **800** | **UV31** | **BUV805** | V31 |  | B31 |  | YG31 |  | R31 |  |
| 805.5-815.5 | 810 | UV32 |  | V32 |  | B32 |  | YG32 |  | R32 |  |
| 815.5-825.3 | 820 | UV33 |  | V33 |  | B33 |  | YG33 |  | R33 |  |
| 825.3-835.1 | 830 | UV34 |  | V34 |  | B34 |  | YG34 |  | R34 |  |
| 835.1-844.7 | 840 | UV35 |  | V35 |  | B35 |  | YG35 |  | R35 |  |

### 2.2 Statement of Ethical Approval, NHP PBMCs and Stimulation

The optimized panel underwent final validation using PBMCs from five adult recycled AGMs, three adult naive olive baboons and two adult naive hybrid olive-yellow baboons. Sex-based analysis was not conducted for this study. Cryopreserved AGM PBMCs (supplemented with 90% FBS and 10% DMSO) were purchased from Bioqual Inc. (Rockville, MD). Due to the limited availability of naive AGMs, PBMCs were sourced from recycled AGMs within a vaccine cohort, 1.5 years following their boost. The vendor has maintained the confidentiality of the vaccine product and immunization regimen. PBMCs from viral-challenged AGMs were not purchased. All experiments involving AGMs were approved by the Institutional Animal Care and Use Committee (IACUC) of Bioqual Inc. (protocol number 22-001). Cryopreserved PBMCs from naive pure and hybrid olive baboon samples, supplemented with 90% FBS and 10% DMSO, were purchased from Southwest National Primate Research Center (SNPRC, San Antonio, TX). The IACUC of SNPRC approved all animal experiments under protocol number 1516PC. At Boston Children’s Hospital, NHP PBMCs were used under the approval of the Institutional Biosafety Committee (Protocol Number: IBC-P00002091). All purchased samples were kept in liquid nitrogen until use.

Cryopreserved NHP PBMCs were thawed in a 37°C water bath. Following thawing, the PBMCs were washed with RPMI 1640 media supplemented with 50 Unit/mL Nuclease (Table 5). Stimulation of resting cells in 1X of T Cell Media (TCM) (22) was achieved using mitogens (see **Section 3.3.2.1: Stimulant Optimization**), followed by incubation in a humidified incubator at 37°C with 5% CO_2_. For the last 5 h of stimulation, Brefeldin A (1:1000) and Monensin (1:1500) (Table 5) were used to inhibit cytokine production. Following this, cells were counted as required before starting the staining process.

### 2.3 Staining Protocol

#### 2.3.1 Materials and Reagents

Table 4 shows the flow antibodies and their titers (final dilution factors) as applied in the 28-color NHP panel. A comprehensive list of all reagents, buffers, plastic consumables, and software/algorithms used in the OMIP’s development can be found in Table 5. Table 6 includes reagents that were used initially but not retained in the optimized panel.

**Table 4.** Optimized 30-parameter/28-color spectral flow panel for NHP species. Deviation of titer of markers in NHPs are highlighted in lime green. For each antibody, the table shows the final dilution (titer), volume per 100 μL test and the specific staining step. Due to downregulation, CXCR5 and 4-1BB staining deviated from the conventional method; these altered steps are marked in magenta.

|  | **Specificity** | **Fluorochrome** | **Clone** | **Vendor** | **Identifier** | **Titer for AGM** | **Titer for Baboon** | **(µL/100 µL) (AGM/Baboon)** | **Staining Step** |
|---|---|---|---|---|---|---|---|---|---|
| 1 | CD3 | APC Vio770 | 10D12 | Miltenyi Biotec | 130-117-695 | 1:40 | 1:20 | 2.5 µL/ 5 µL | Surface |
| 2 | CD4 | BV786 | L200 | Waters Biosciences | 563914 | 1:80 | 1:40 | 1.25 µL/ 2.5 µL | Surface |
| 3 | CD8 | RB545 | SK1 | Waters Biosciences | 756508 | 1:40 | 1:160 | 2.5 µL/ 0.625 µL | Surface |
| 4 | CD11b | RB613 | ICRF44 | Waters Biosciences | 571258 | 1:40 | 1:40 | 2.5 µL for both species | Surface |
| 5 | CD11c | BV650 | S-HCL-3 | Waters Biosciences | 744437 | 1:40 | 1:40 | 2.5 µL for both species | Surface |
| 6 | CD16 | BUV496 | 3G8 | Waters Biosciences | 612944 | 1:80 | 1:40 | 1.25 µL/ 2.5 µL | Surface |
| 7 | CD20 | RB780 | 2H7 | Waters Biosciences | 569120 | 1:20 | 1:20 | 5 µL for both species | Surface |
| 8 | CD25 | PE Fire 640 | M-A251 | BioLegend | 356148 | 1:40 | 1:40 | 2.5 µL for both species | Surface |
| 9 | CD45 | BV510 | D058-1283 | Waters Biosciences | 563530 | 1:40 | 1:40 | 2.5 µL for both species | Surface |
| 10 | CD45RA | BUV805 | 5H9 | Waters Biosciences | 742052 | 1:80 | 1:80 | 1.25 µL for both species | Surface |
| 11 | CD56 | BUV563 | NCAM16.2 | Waters Biosciences | 612928 | 1:160 | 1:320 | 0.625 µL/ 0.3125 µL | Surface |
| 12 | CD69 | RY610 | FN50 | Waters Biosciences | 571167 | 1:40 | 1:40 | 2.5 µL for both species | Surface |
| 13 | CD95 | R718 | DX2 | Waters Biosciences | 569572 | 1:160 | 1:160 | 0.625 µL for both species | Surface |
| 14 | CD123 | RY703 | 7G3 | Waters Biosciences | 571445 | 1:40 | 1:40 | 2.5 µL for both species | Surface |
| 15 | CD134/OX-40 | BUV615 | L106 | Waters Biosciences | 751320 | 1:80 | 1:80 | 1.25 µL for both species | Surface |
| 16 | CD137/4-1BB | BV711 | 4B4-1 | Waters Biosciences | 569689 | 1:40 | 1:40 | 2.5 µL for both species | Pre-cytokine blockage |
| 17 | CD159a/NKG2a | APC | Z199 | Beckman Coulter | A60797 | 1:40 | 1:40 | 2.5 µL for both species | Surface |
| 18 | CD185/CXCR5 | FITC | MU5UBEE | Thermo Fisher | 11-9185-42 | 1:160 | 1:160 | 0.625 µL for both species | Pre-stimulation |
| 19 | CD279/PD-1 | BV605 | EH12.1 | Waters Biosciences | 563245 | 1:40 | 1:40 | 2.5 µL for both species | Surface |
| 20 | FoxP3 | AF647 | 206D | BioLegend | 320114 | 1:20 | 1:20 | 5 µL for both species | Nuclear* |
| 21 | HLA-DR | BV570 | L243 | BioLegend | 307638 | 1:40 | 1:40 | 2.5 µL for both species | Surface |
| 22 | IFNγ | RB705 | B27 | Waters Biosciences | 570247 | 1:20 | 1:20 | 5 µL for both species | Nuclear* |
| 23 | IL-2 | BUV737 | MQ1-17H12 | Waters Biosciences | 612836 | 1:40 | 1:40 | 2.5 µL for both species | Nuclear |
| 24 | IL-4 | PE | 8D4-8 | Thermo Fisher | 12-7049-42 | 1:40 | 1:40 | 2.5 µL for both species | Nuclear |
| 25 | IL-5 | PE | JES1-39D10 | BioLegend | 500904 | 1:40 | 1:80 | 2.5 µL/ 1.25 µL | Nuclear |
| 26 | IL-13 | PE | JES10-5A2 | BioLegend | 501903 | 1:40 | 1:40 | 2.5 µL for both species | Nuclear |
| 27 | IL-17A | BV421 | N49-653 | Waters Biosciences | 562933 | 1:20 | 1:20 | 5 µL for both species | Nuclear |
| 28 | TCRγδ | RB744 | B1 | Waters Biosciences | 758180 | 1:20 | 1:20 | 5 µL for both species | Surface |
| 29 | TNF | PE Cy7 | MAb11 | Waters Biosciences | 557647 | 1:20 | 1:20 | 5 µL for both species | Nuclear |
| 30 | Viability | Zombie NIR | --- | BioLegend | 423106 | 1:1000 | 1:1000 | ---- | Post-Fc blockage |

\* Antibody validation involved both ICS and nuclear buffer. However, the final panel specifically used nuclear buffer because of the need to include FoxP3.

**Table 5.**
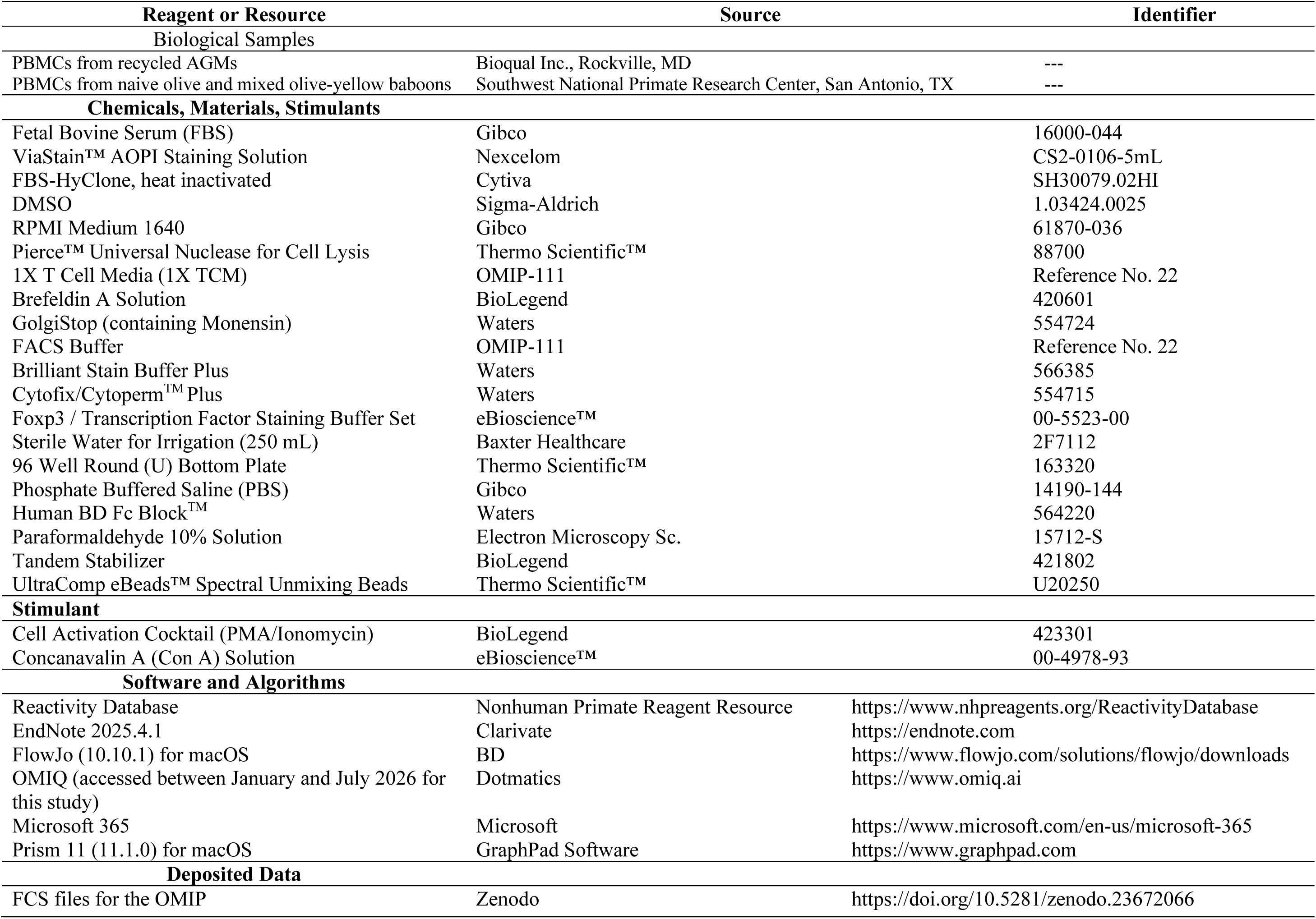
Biological samples, reagents, stimulants and algorithms.

**Table 6.**
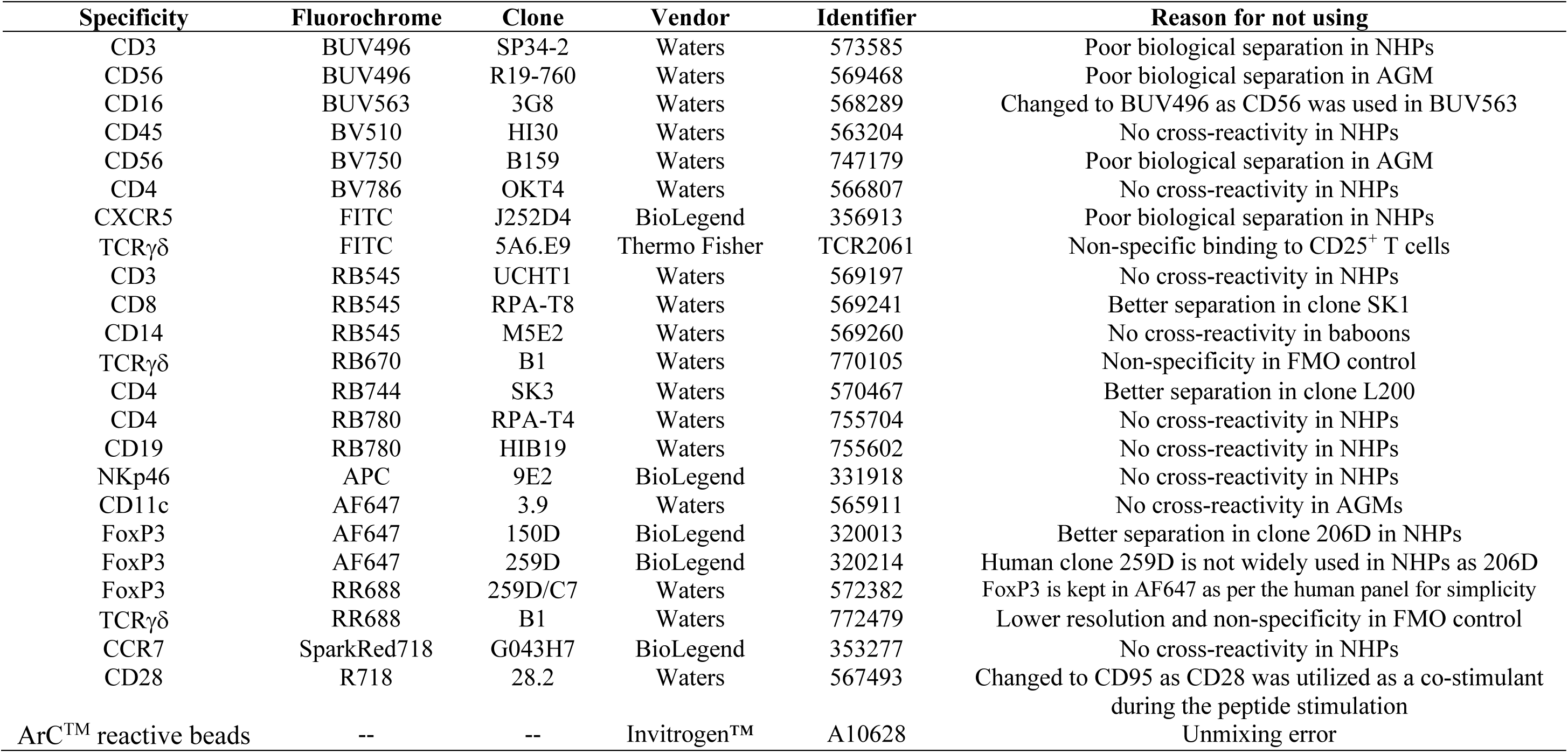
Commercial reagents were evaluated but excluded from the final optimized panel.

#### 2.3.2 Antibody preparation, Fc blocking and Viability staining

#### 2.3.3 Surface Antibody Staining

#### 2.3.4 Intracellular or Intranuclear Staining

#### 2.3.5 Fixation and Storage

#### 2.3.6 Cell Analysis using ID7000™

The above sections (from **2.3.2** to **2.3.6**) adopt the methodology outlined in OMIP-XXX-A (3). To prevent repetitive descriptions, the procedures for **antibody preparation**, **Fc blocking**, **viability** and **surface staining**, subsequent **intracellular** or **nuclear staining**, **fixation**, and **cell acquisition** via ID7000™ are directly referenced from OMIP-XXX-A (3).

## 3 Panel Design and Optimization

### 3.1 Panel Design

The primary objective of this OMIP’s development was to establish a broad-spectral panel applicable to NHPs, particularly AGMs and baboons. This was driven by their utility in current vaccine efficacy studies in AGMs and infant baboon challenge models for pertussis pathogenesis. We identified antibodies with cross-species reactivity between NHP and human by searching the NHP “Reactivity Database” (21) and the OMIP database (20). We assigned these antibodies to respective fluorochromes in accordance with the recently published panel development guidelines (27–29). To mitigate unmixing errors, we leveraged Sony Biotechnology’s PDT for spectral overlap prediction, as depicted in Figure 2. Our aim was to establish OMIP-XXX-A (3) as an NHP equivalent, given OMIP-XXX-B’s human-specific reagents. Thus, human PBMCs were included as a positive control in each NHP panel run (iteration, Table 7). The antibodies for the backbone panel in **Ultraviolet (355 nm)**, **Violet (405 nm)**, **Blue (488 nm)**, **Yellow-Green (561 nm)**, and **Red (637 nm)** spectra were selected as described in **Section 3.1.2: Fluorochrome Assignment** of OMIP-XXX-A (3), with the exception noted below.

**Figure 2:**
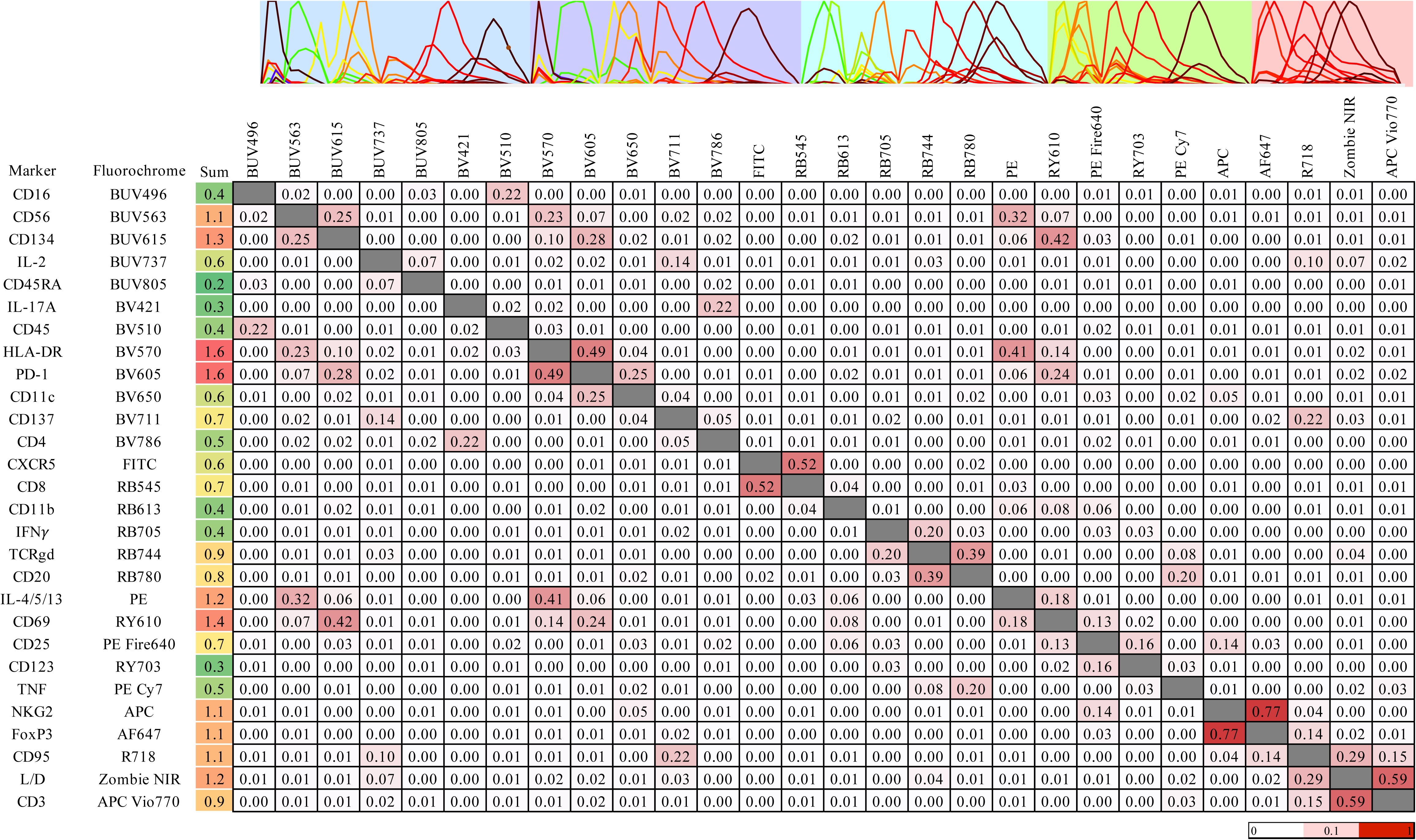
The figure shows the Similarity Index Matrix (SIM) for the optimized 28-color NHP panel on the ID7000™ spectral cell analyzer. The SIM, generated by Sony Biotechnology Inc.’s Panel Development Tool (PDT), quantifies the spectral similarity between fluorophores. A score from 0 to 1 indicates similarity, with higher value (shown in red on the heatmap) meaning greater similarity. Above the SIM, the spectral distribution of all fluorochromes is displayed, based on reference spectra and the instrument’s laser configuration to ensure precise spectral unmixing. The background color indicate the excitation laser lines: 355 nm (pale blue), 405 nm (pale purple), 488 nm (ice), 561 nm (lime green) and 637 nm (salmon). The figure was generated by Sony Biotechnology’s PDT.

**Table 7.** Iterative (1-26) optimization of the NHP panel. For each phase, the human panel variant was run in parallel using human PBMCs. Iterations highlighted in red (25 & 26) were evaluated independently.

| Detector | Fluorochrome | Iteration 1 | Iteration 2 | Iteration 3 | Iteration 4 | Iteration 5 | Iteration 6 | Iteration 7 | Iteration 8 | Iteration 9 | Iteration 10 | Iteration 11 | Iteration 12 | Iteration 13 | Iteration 14 |
|---|---|---|---|---|---|---|---|---|---|---|---|---|---|---|---|
| UV1 | BUV395 | CD8 | CD8 | CD8 | CD8 | CD8 |  |  |  |  |  |  |  |  |  |
| UV3 | BUV496 | CD3 |  | CD3 | CD3 |  |  |  |  |  |  |  |  |  | CD56 |
| UV9 | BUV563 |  |  |  |  | CD16 |  |  | CD16 | CD16 | CD16 | CD16 | CD16 | CD16 | CD16 |
| UV31 | BUV805 |  |  |  |  | CD45RA |  |  |  |  |  |  |  |  |  |
| V5 | BV510 | CD45 | CD45 | CD45 | CD45 | CD45 | CD45 | CD45 | CD45 | CD45 | CD45 | CD45 | CD45 | CD45 | CD45 |
| V10 | BV570 |  |  |  |  | HLA-DR |  |  | HLA-DR | HLA-DR | HLA-DR | HLA-DR | HLA-DR | HLA-DR | HLA-DR |
| V13 | BV605 | PD-1 | PD-1 | PD-1 | PD-1 |  |  |  |  |  |  |  | PD-1 |  |  |
| V17 | BV650 |  |  |  |  | CD11c |  |  |  |  |  | CD11c |  |  |  |
| V26 | BV750 |  |  |  |  |  |  |  |  |  |  |  |  | CD56 |  |
| V29 | BV786 |  | CD4 |  |  | CD4 | CD4 |  |  |  |  |  | CD4 |  |  |
| B4 | FITC | CXCR5 | CXCR5 | CXCR5 | CXCR5 |  |  |  |  |  |  |  | CXCR5 |  |  |
| B8 | RB545 |  |  |  |  |  | CD8 |  |  | CD14 |  |  | CD8 |  |  |
| B14 | RB613 |  |  |  |  |  |  |  | CD11b |  | CD11b | CD11b |  |  |  |
| B26 | RB744 | TCRγδ | TCRγδ | CD4 |  | TCRγδ |  |  |  |  |  |  |  |  |  |
| B29 | RB780 | CD20 | CD20 | CD20 | CD4 | CD20 | CD20 | CD19 | CD20 | CD20 | CD20 | CD20 | CD20 | CD20 | CD20 |
| YG11 | RY586 |  |  |  | CD20 |  |  |  |  |  |  |  |  |  |  |
| YG16 | PE Fire640 | CD25 | CD25 | CD25 | CD25 | CD25 |  |  |  |  |  |  |  |  |  |
| YG22 | RY703 |  |  |  |  |  |  |  |  |  | CD123 | CD123 |  |  |  |
| R18 | APC |  |  |  |  | NKG2 |  |  |  |  |  |  | NKG2 | NKG2 | NKG2 |
| R19 | AF647 |  |  |  |  |  |  |  |  |  | CD11c |  |  |  |  |
| R23 | R718 |  |  |  |  | CD28 |  |  |  |  |  |  |  |  |  |
| R26 | Zombie NIR | Viability | Viability | Viability | Viability | Viability | Viability | Viability | Viability | Viability | Viability | Viability | Viability | Viability | Viability |
| R28 | APC H7 | CD4 |  |  |  |  |  |  |  |  |  |  |  |  |  |
| R28 | APC Vio770 |  | CD3 |  |  | CD3 | CD3 | CD3 | CD3 | CD3 | CD3 | CD3 | CD3 | CD3 | CD3 |
| UV3 | BUV496 | CD16 |  |  |  |  |  |  | CD56 |  |  | CD16 | CD16 | CD16 |  |
| UV9 | BUV563 | CD56 |  |  |  |  | CD16 | CD16 | CD16 |  |  | CD56 | CD56 | CD56 |  |
| UV14 | BUV615 |  |  |  |  |  |  |  | CD134 |  |  | CD134 |  | CD134 |  |
| UV25 | BUV737 |  |  |  |  |  |  |  | IL-2 |  |  |  | IL-2 | IL-2 |  |
| UV31 | BUV805 |  |  |  |  |  |  |  | CD45RA | CD45RA | CD45RA | CD45RA | CD45RA | CD45RA |  |
| V1 | BV421 |  |  |  |  |  |  |  | IL-17 |  |  |  | IL-17A | IL-17A |  |
| V5 | BV510 | CD45 | CD45 | CD45 | CD45 | CD45 | CD45 | CD45 | CD45 | CD45 | CD45 | CD45 | CD45 | CD45 |  |
| V10 | BV570 | HLA-DR |  |  |  |  | HLA-DR | HLA-DR | HLA-DR |  |  | HLA-DR | HLA-DR | HLA-DR |  |
| V13 | BV605 |  |  |  |  |  |  |  | PD-1 |  |  | PD-1 |  | PD-1 |  |
| V17 | BV650 |  |  |  |  |  |  |  | CD11c |  |  | CD11c |  | CD11c |  |
| V22 | BV711 |  |  |  |  |  |  |  | CD137 |  |  | CD137 |  | CD137 |  |
| V29 | BV786 |  | CD4 | CD4 | CD4 | CD4 | CD4 | CD4 | CD4 | CD4 | CD4 | CD4 | CD4 | CD4 |  |
| B4 | FITC |  |  | TCRγδ |  |  |  |  | CXCR5 |  |  | CXCR5 |  | CXCR5 |  |
| B8 | RB545 |  | CD8 | CD8 | CD8 | CD8 | CD8 | CD8 | CD8 | CD8 | CD8 | CD8 | CD8 | CD8 |  |
| B14 | RB613 |  |  |  |  |  |  |  | CD11b |  |  | CD11b |  | CD11b |  |
| B19 | RB670 |  |  |  | TCRγδ |  |  |  |  |  |  |  |  |  |  |
| B22 | RB705 |  |  |  |  |  |  |  | IFNγ |  |  |  | IFNγ | IFNγ |  |
| B26 | RB744 |  | TCRγδ |  |  |  |  |  | TCRγδ | TCRγδ | TCRγδ |  |  | TCRγδ |  |
| B29 | RB780 | CD20 | CD20 | CD20 | CD20 | CD20 | CD20 | CD20 | CD20 | CD20 | CD20 | CD20 | CD20 | CD20 |  |
| YG11 | PE |  |  |  |  |  |  |  | IL-4/5/13 |  |  |  | IL-4/5/13 | IL-4/5/13 |  |
| YG13 | RY610 |  |  |  |  |  |  |  | CD69 |  |  | CD69 | CD69 | CD69 |  |
| YG16 | PE Fire640 |  | CD25 | CD25 | CD25 | CD25 | CD25 | CD25 | CD25 |  |  | CD25 |  | CD25 |  |
| YG22 | RY703 |  |  |  |  |  |  |  | CD123 |  |  | CD123 |  | CD123 |  |
| YG29 | PE Cy7 |  |  |  |  |  |  |  | TNF |  |  |  | TNF | TNF |  |
| R18 | APC | NKG2 |  |  |  |  | NKG2 | NKG2 | NKG2 |  |  | NKG2 | NKG2 | NKG2 |  |
| R19 | AF647 |  |  |  |  |  | FoxP3 |  |  |  |  |  |  | FoxP3 |  |
| R20 | RR688 |  |  |  |  | TCRγδ |  | FoxP3 |  |  |  |  |  |  |  |
| R23 | SparkRed718 |  |  |  |  |  |  |  | CCR7 |  |  |  |  |  |  |
| R23 | R718 |  |  |  |  |  |  |  |  | CD28 | CD95 | CD95 | CD95 | CD95 |  |
| R26 | Zombie NIR | Viability | Viability | Viability | Viability | Viability | Viability | Viability | Viability | Viability | Viability | Viability | Viability | Viability |  |
| R28 | APC Vio770 | CD3 | CD3 | CD3 | CD3 | CD3 | CD3 | CD3 | CD3 | CD3 | CD3 | CD3 | CD3 | CD3 |  |

For NHPs, CD3 clone 10D12 outperformed SP34-2, so we kept it on a dim fluorochrome APC Vio770. In the human panel, we used CD8 (RPA-T8) in BUV395. We retained CD8 clone SK1 on moderate fluorochrome RB545 due to its superior resolution in AGM. For further details on these specifications, please refer to **Section 3.2.2: Panel Iterations and FMO Controls**, below. Consistent with Figure 2, Figure 3 also demonstrates minimal spectral spillover.

**Figure 3:**
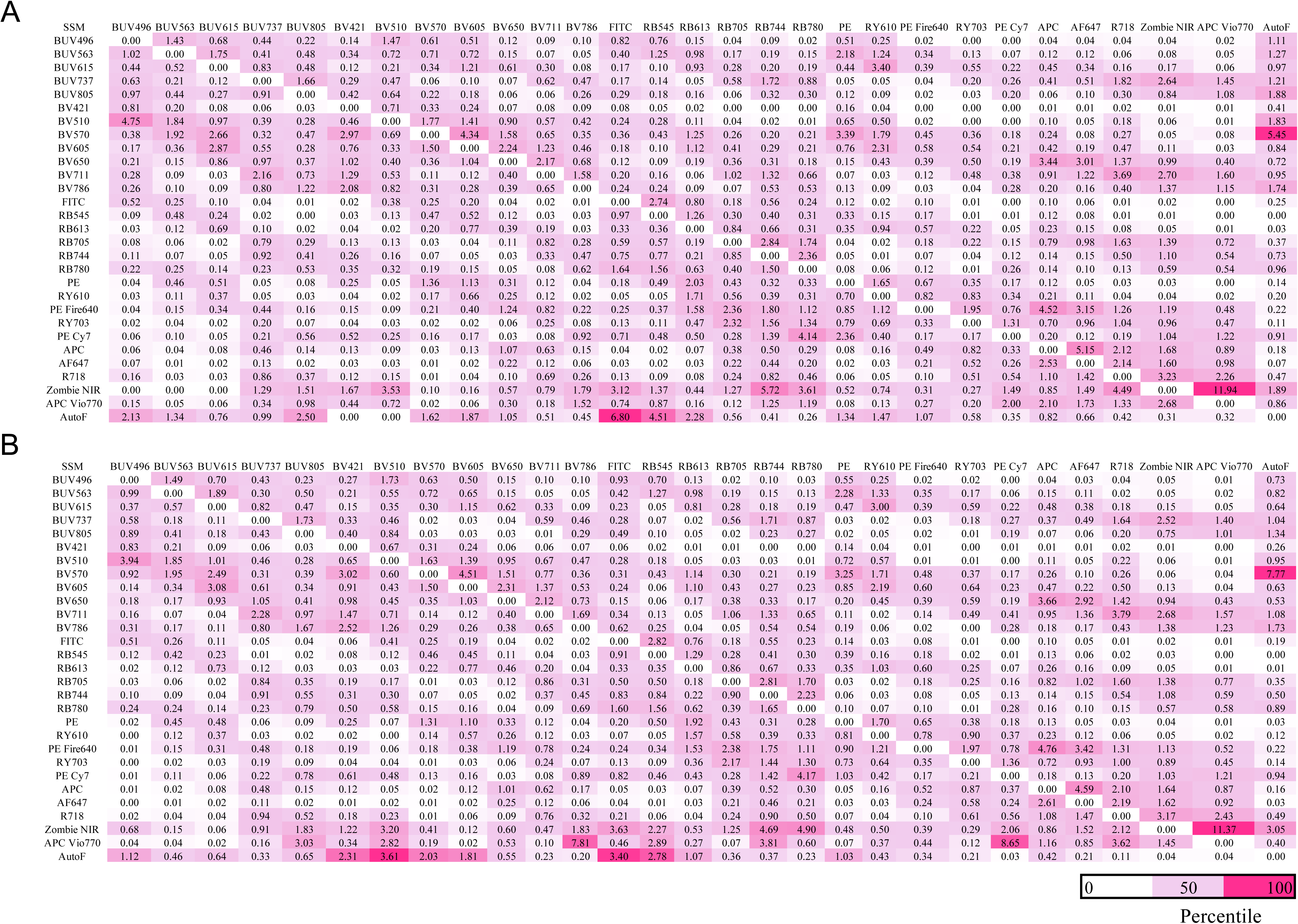
The Spillover Spreading Matrix (SSM) for both the optimized 28-color (A) AGM and (B) baboon panels, acquired on the Sony ID7000™ spectral cell analyzer. This SSM was generated using FlowJo’s (10.10.1) AutoSpill algorithm, from single-color controls that included UltraComp eBeads™ Spectral Unmixing Beads and NHP-specific cellular controls for BV570 and Zombie NIR to reduce unmixing errors. Unstained cells (NHP specific) were used as an autofluorescence (AutoF) parameter during unmixing of the data. Each cell value represents the impact of spillover spread between individual pairs of fluorochromes. Spillover values are color-coded by percentile, with light plum indicating the 50th percentile and strawberry representing high values.

### 3.2 Panel Optimization

#### 3.2.1 Antibody Titrations using combinatorial titration method

We implemented the Combinatorial Antibody Titrations (CAT) method (30) to simultaneously titrate multiple antibodies, thereby conserving NHP PBMCs as described in OMIP-XXX-A, section 3.2.1 (3). The strategy for executing the CAT method is illustrated in Figure 4. We consistently applied a similarly index cut off of ≤ 0.1 for any two or more chosen test fluorochromes within each CAT group. To address potential unmixing errors during antibody titration for both AGMs and olive baboons, we tested a total of 13 combinations. Our evaluation included further division of Groups 2 and 8 into subgroups (Figure 4). Specifically, Groups 2.1 and 8.1 were dedicated to AGMs, while 2.2 and 8.2 focused on olive baboons, driven by the need to evaluate CD45RA and CD8 clones (further explained in **Section 3.2.2**). For olive baboons, CD8 (SK1) RB545 was titrated as a standalone antibody with viability marker, and its SIM is not shown in Figure 4 to prevent redundancy with Group 2.1. Similarly, IL-13 PE underwent solo titration, incorporating viability and CD3 as a lineage marker, mirroring the approach used for Group 3 or 4 (Figure 4). Our overall strategy involved titrating all antibodies using cryopreserved PBMCs, either in their steady state or after mitogen stimulation and overnight resting, consistent with the methodology described in OMIP-XXX-A (3). The effectiveness of the CAT method was re-validated, as illustrated in Figure 4B-E. This re-validation utilized PBMCs from both AGMs (Figures 4B-D) and olive baboons (Figures 4E), specifically targeting the ICS marker IFNγ. Optimal titration was determined by a comprehensive assessment process, which involved visualizing the concatenated bivariate plots for each marker (Figures 4B-C, 5.1 & 5.2) alongside its corresponding Stain Index curve (Figures 4D-E, 6.1 & 6.2), a methodology detailed in OMIP-XXX-A (3). Within these visual representations, red rectangles in the bivariate plots (Figures 4B-C, 5.1 & 5.2) and dark blue rectangles on the Stain Index curves (Figures 4D-E, 6.1 & 6.2) denote the selected optimal titers. Table 4 also illustrates the titration between NHP species, with the deviation between species being highlighted in lime green.

**Figure 4:**
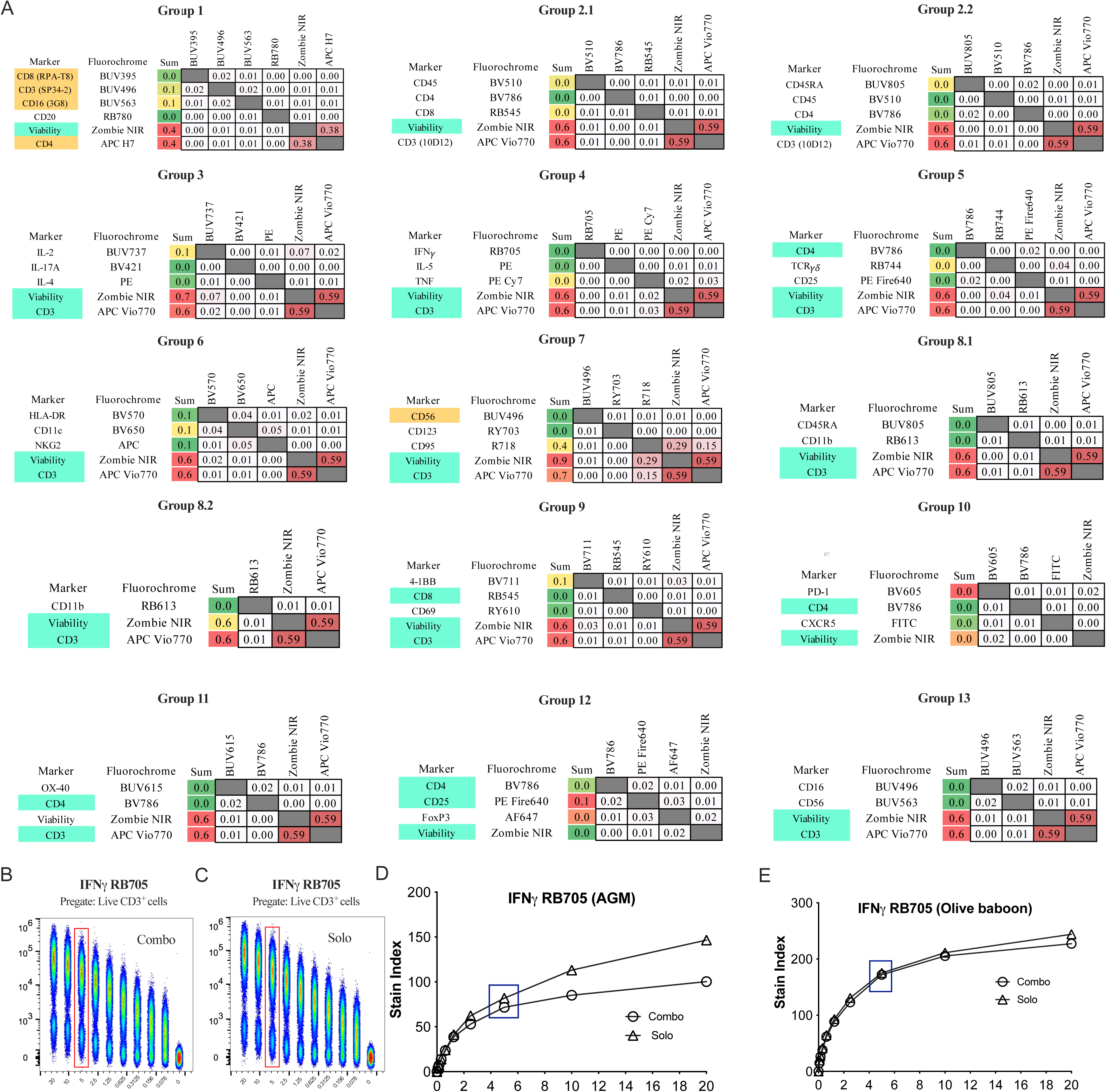
(A) We used a “combinatorial antibody titrations” strategy with PDT (Sony Biotechnology Inc.) to predict the lowest similarity indices on the SIM, thereby reducing unmixing errors. This strategy applied to PBMCs from both NHP species, except for groups 2 and 8. Groups 2.1 and 8.1 were designed for AGMs, while groups 2.2 and 8.2 were for olive baboons. We set the cut-off value for similarity score at less than 0.1 between fluorochromes. In SIMs, we illustrated lineage markers in aquamarine, a strategy developed to titrate rare markers. Markers in orange were used during panel development but not included (**Table 6**) in the final panel. We further validated the combinatorial method anonymously using the ICS marker IFNγ. Visual comparisons of concatenated representations showed (B) the combinatorial and (C) traditional methods in AGM PBMCs. The Stain Index curve of IFNγ demonstrated consistency between the two distinct titration methods in (D) AGMs and (E) olive baboons. The rectangular box in dark blue indicated the chosen titer.

**Figure 5.1:**
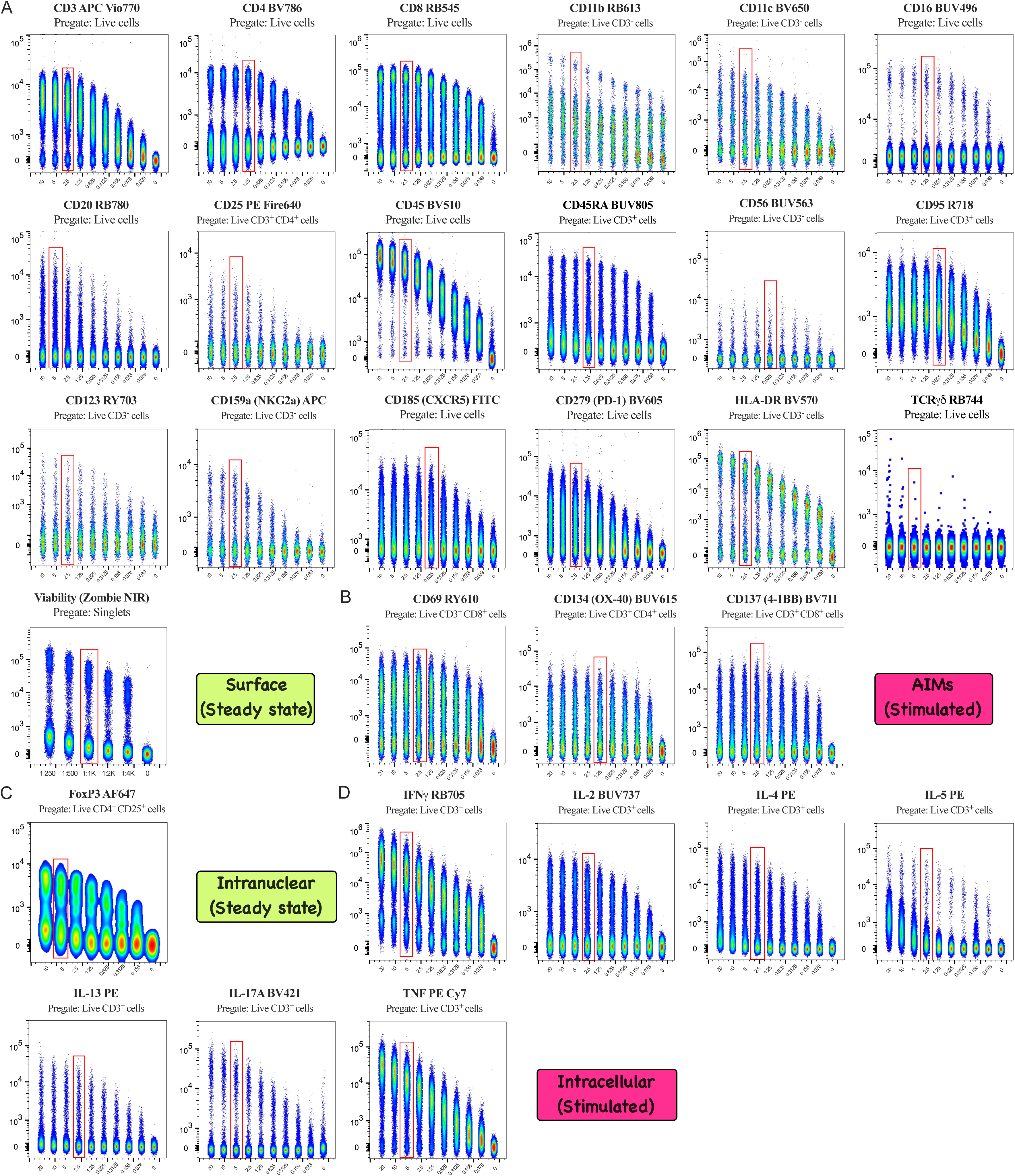
Antibody concentrations were optimized through titrations using cryopreserved, overnight-rested AGM PBMCs. Details of the methodology can be found in Figure 5.2.

**Figure 5.2:**
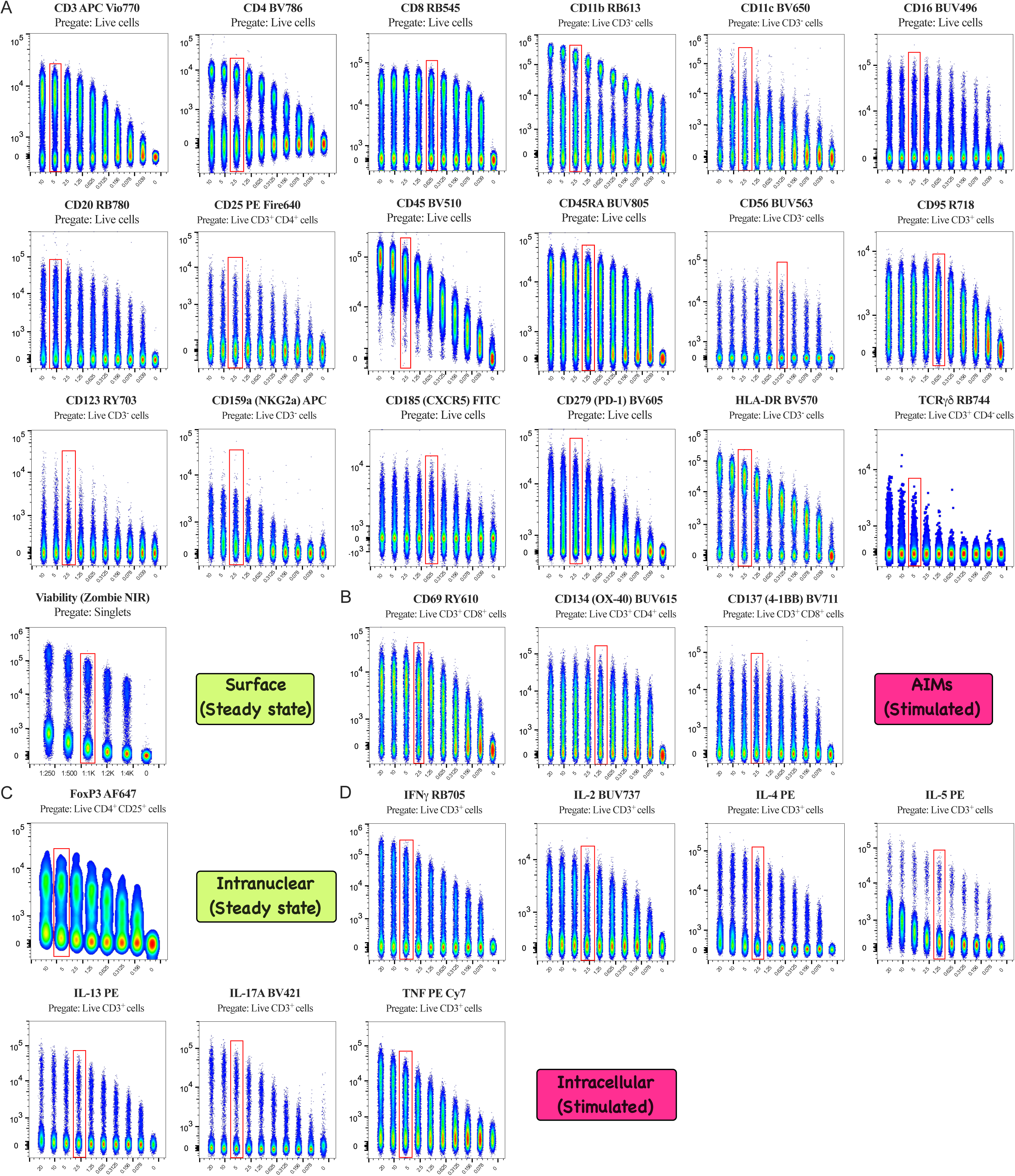
Optimal concentration of antibodies is defined by titrations using overnight rested cryopreserved PBMCs from Olive baboons. Serial fold-dilutions (on the X-axis) of individual antibodies were initiated at either 20 µL/100 µL or 10 µL/100 µL, with the exception of L/D Zombie NIR, which began at a 1:250 dilution. For (A) surface antibodies, titrations were conducted on steady-state NHP PBMCs. In the case of (B) AIMs, cells underwent a 20 h stimulation with Concanavalin A prior to titration, while for (D) intracellular cytokines, cells were stimulated for 5 h with a mitogen before the titration process. FCS files were processed by gating out debris (FSC/SSC) and doublets, followed by gating on live cells or specific indicated subsets, and then concatenated for representation. The chosen titer, highlighted by red rectangles, was selected from the Stain Index curve (Figure 6), ensuring distinct separation between positive and negative populations with minimal background staining. All selected antibody titrations are also documented in **Table 4**, expressed as either µL/100 µL or as a dilution factor.

**Figure 6.1:**
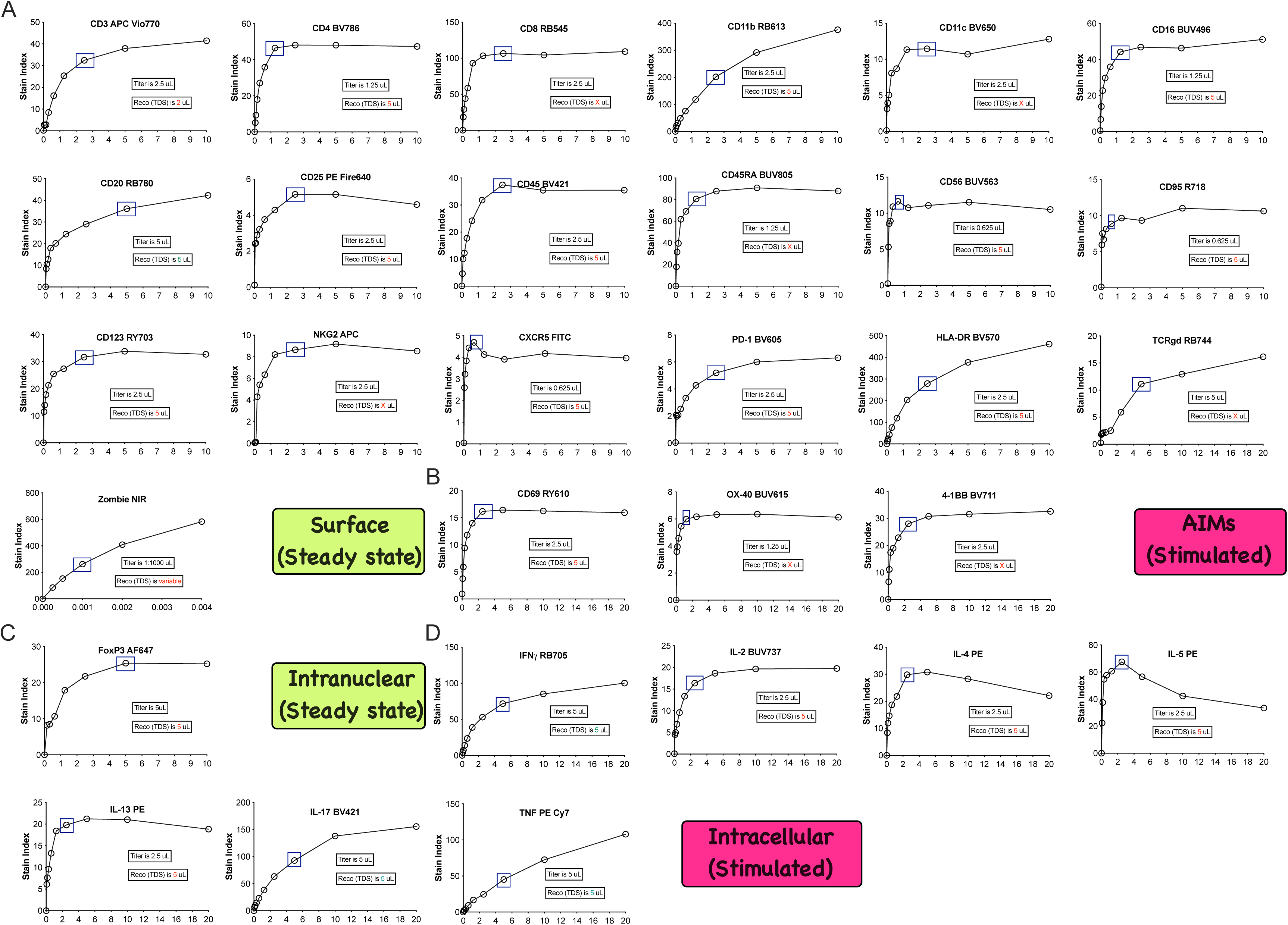
The Stain Index curve for the selected 30 markers in the optimized AGM panel is shown. A dark blue rectangular box marks the chosen titration point where a plateau is observed. Each plot also indicates the chosen titers. Furthermore, the recommended (reco) titration is presented, based on the technical data sheet (TDS) of each reagent, to highlight any discrepancies. All (A) surface, (B) AIMs, (C) intranuclear, and (D) intracellular cytokines were titrated using AGM PBMCs under either steady-state or stimulated conditions, as further detailed in Figure 5.1.

**Figure 6.2:**
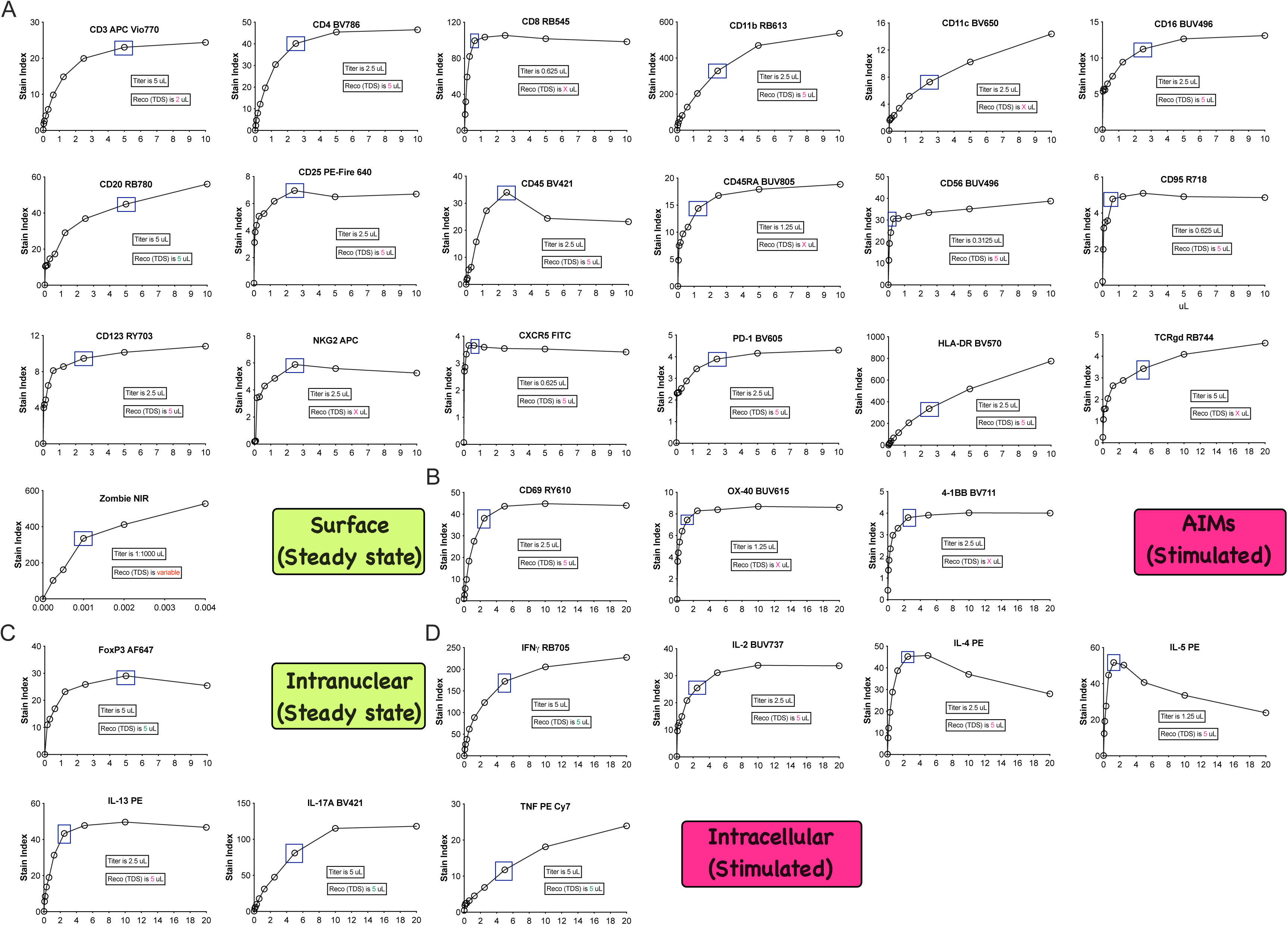
The Stain Index curve for the 30 selected markers of the optimized olive baboon panel is illustrated. A dark blue rectangular box points out the chosen titration point where a plateau is evident. The selected titers are further marked within each plot. Additionally, the recommended (reco) titration is provided, derived from the technical data sheet (TDS) of each reagent to reveal discrepancies. The titrations encompassed (A) surface, (B) AIMs, (C) intranuclear, and (D) intracellular cytokines, utilizing olive baboons PBMCs in either steady-state or stimulated conditions, as detailed in Figure 5.2.

#### 3.2.2 Panel Iterations and FMO Controls

The initial panel was conceptualized using Sony Biotechnology’s PDT alongside the NHP Reactivity Database (21) to predict cross-species antibody reactivity (See **Section 3.1**). Iterative panel optimization was conducted on co-processed PBMCs across all targeted species (Table 7). During T-cell lineage identification, CD3 clone 10D12 out-performed clone SP34-2, providing superior CD3^+^ population separation in both AGM and baboons (Figures 7.1A & 7.1B) while preserving non-B/T cell resolution in AGMs. Consequently, clone 10D12 was incorporated using the dim fluorochrome APC Vio770. Cross-species lineage profiling revealed that 10D12 lacked human reactivity (Figure 7.1B), whereas the human-specific CD3 clone UCHT1 failed to label NHP CD3^+^ T cells (Figure 7.1C). For CD8 lineage marker evaluation, clone SK1 (Figures 7.1E) yielded distinctively better population resolution in AGMs than clone RPA-T8 (Figures 7.1D) and was assigned to the moderate fluorochrome RB545. Lastly, of four CD4 clones tested (Figures 7.1G-J), clone L200 was selected for downstream human and NHP analyses given its enhanced performance in NHPs, consistent with its prior validation in baboons (25).

**Figure 7.1:**
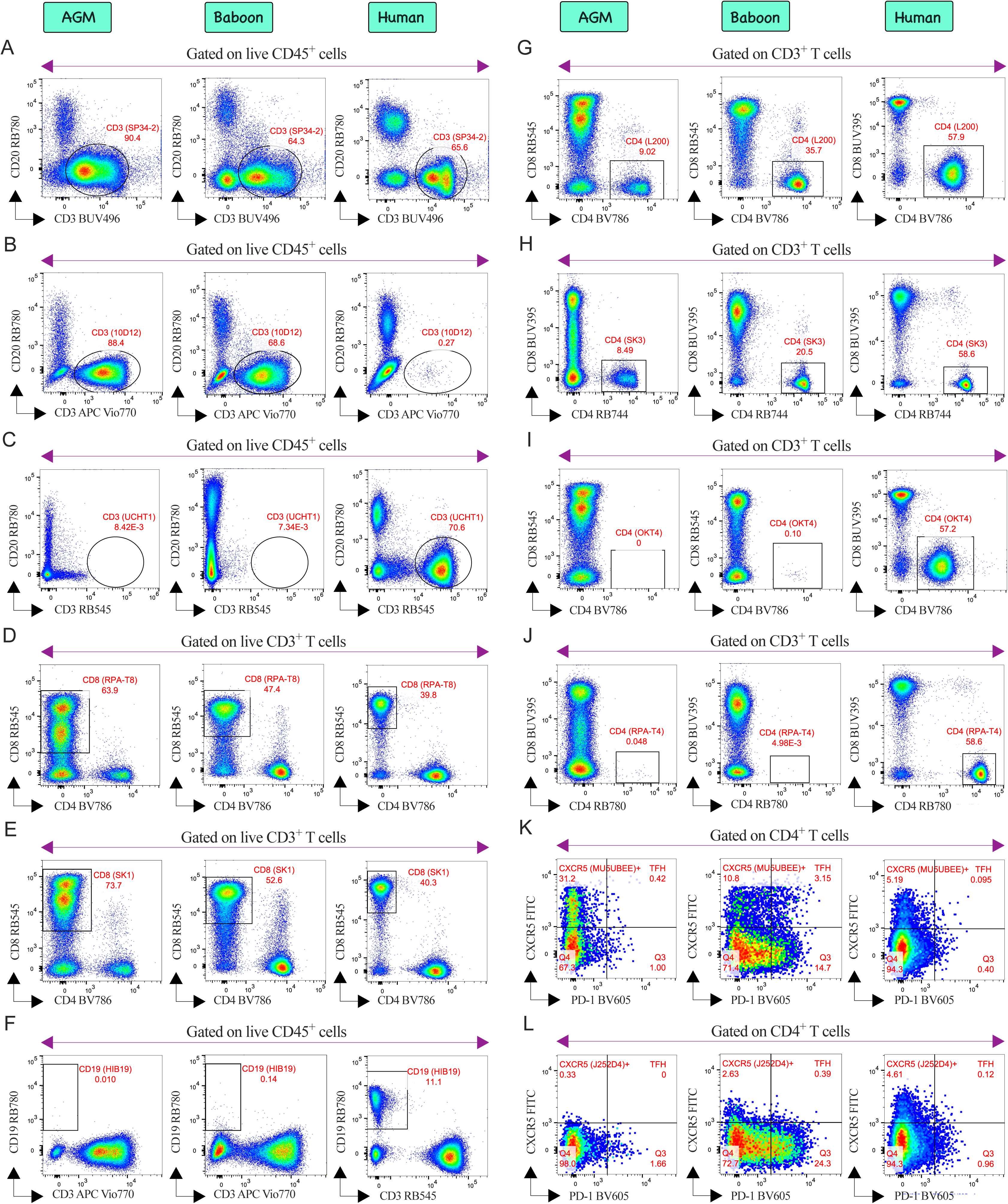
Evaluation of various clones of (A-C) CD3, (D, E) CD8, (F) CD19, (G-J) CD4, and (K, L) CXCR5 antibodies across NHPs and humans. Based on their performance, specific clones were selected for further analysis. Each column represents a distinct species, as indicated at the top.

B cells were identified using CD20 alone (Figures 1A & 1H), because CD19 clone HIB19 lacks NHP cross-reactivity (Figure 7.1F). For TFH cell identification, CXCR5 clone MU5UBEE (Figure 7.1K), previously validated in OMIP-75 (2), outperformed clone J252D4 in AGMs, olive baboons, and humans (Figure 7.1L). Finally, re-evaluating pan-leukocyte CD45 clones D058-1283 (NHP-specific; Figure 7.2A) and HI30 (human-specific; Figure 7.2B) confirmed clear species-specific bias. To phenotype monocyte stages, we evaluated the standard CD14 and CD16 combination (Figure 7.2C). Because CD14 clone M5E2 lacked reactivity in baboons (Figure 7.2C), we instead adopted a CD11b/CD16 profiling strategy, originally reported by the Stanford group (31), to consistently define monocyte stages across all species (Figure 7.2D). Contrary to their findings, we observed no CD11b non-reactivity in AGMs (Figure 7.2D). For CD11c, clone S-HCL-3 was selected (Figure 7.2F) after clone 3.9 failed in AGMs (Figure 7.2E), which likely explains the Stanford group’s inability (31) to detect myeloid dendritic cells (mDCs) in this species. Human NK cells were characterized using NKp46 and CD16 (Figure 7.2G) per OMIP-70 (32); however, NKp46 lacked cross-reactivity in NHPs (Figure 7.2G). To bypass this, NHP NK cells were identified using NKG2a clone Z199 (Figure 7.2H), a strategy previously established in AGMs (33) and in rhesus macaques (34–37). Among three CD56 clones evaluated (Figure 7.2I-K), only NCAM16.2 (Figure 7.2K) exhibited the strongest cross-species reactivity. Furthermore, pairing CD56 BUV563 with CD16 BUV496 (Figure 7.2L) provided improved resolution of early NK cells across all species compared to the reciprocal CD56 BUV496/CD16 BUV563 combination (Figure 7.2K) -a strategy consistent with previously validated OMIPs (38–41). The differential expression of early NK cells in recycled AGMs (Figures 1B, 7.2L & 8D) likely reflects high viremia (6), which drives the expansion or recruitment of mature effector NK cells (7).

**Figure 7.2:**
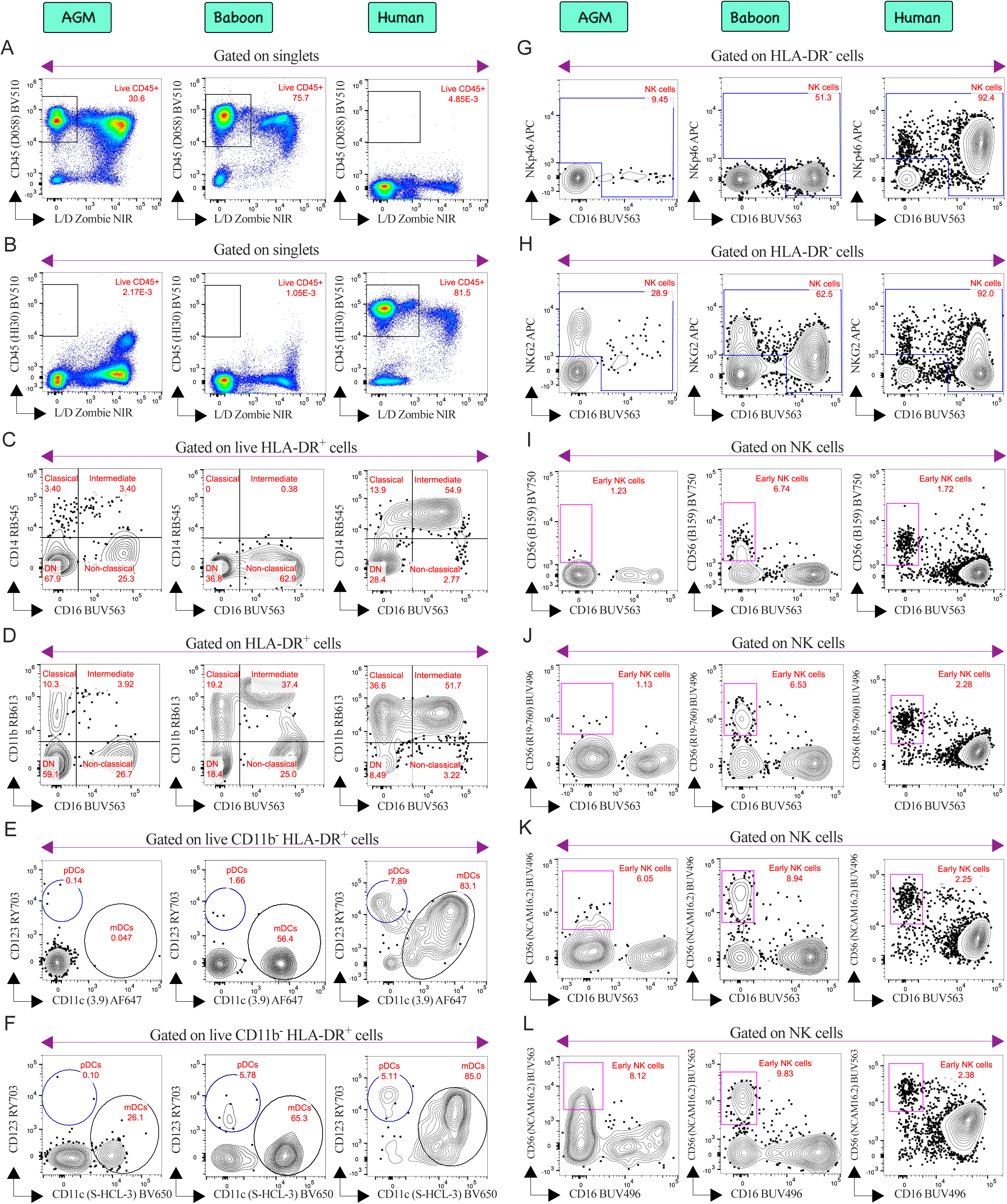
Evaluation of various clones of (A-B) CD45, (E, F) CD11c, (I-L) CD56 antibodies across NHPs and humans. Based on their performance, specific clones were selected for further analysis. Furthermore, the combination of (D) CD11b and CD16 was employed to depict monocytes phages, as (C) CD14 proved ineffective in Baboons. Similarly, (G) NKp46 in humans and (H) NKG2 clone in NHPs demonstrated efficiency in capturing NK cells. Each column represents a distinct species, as indicated at the top.

According to NHP cross-reactivity databases (21), clones 5A6.E9 and B1 are recommended to phenotype TCRγδ^+^ T cells in AGMs and baboons, respectively. While 5A6.E9 displayed broad species reactivity (Figure 7.3A), it displayed unexpected non-specific binding to CD3^+^ CD25^+^ T cells (Figure 7.3B). Investigators should exercise caution when using 5A6.E9 due to this off-target reactivity. Conversely, clone B1 eliminated this off-target artifact (Figure 7.3C) and performed well in human and baboon samples (Figure 7.3D), but detected lower TCRγδ cell frequencies in AGMs. To optimize B1 performance, PBMCs from three AGMs were evaluated using iterations 16, 18, and 19 (Table 7) to compare TCRγδ RB744 against RB670 and RR688 (Figure 7.4). Although RB670 yielded relatively higher staining intensity across subjects (Figure 7.4D), RB744 was selected due to its superior specificity during FMO assessment (Figure 7.4E). Notably, inter-individual variability in TCRγδ cell frequency among AGMs likely reflects their non-naive status as previously exposed, recycled animals.

**Figure 7.3:**
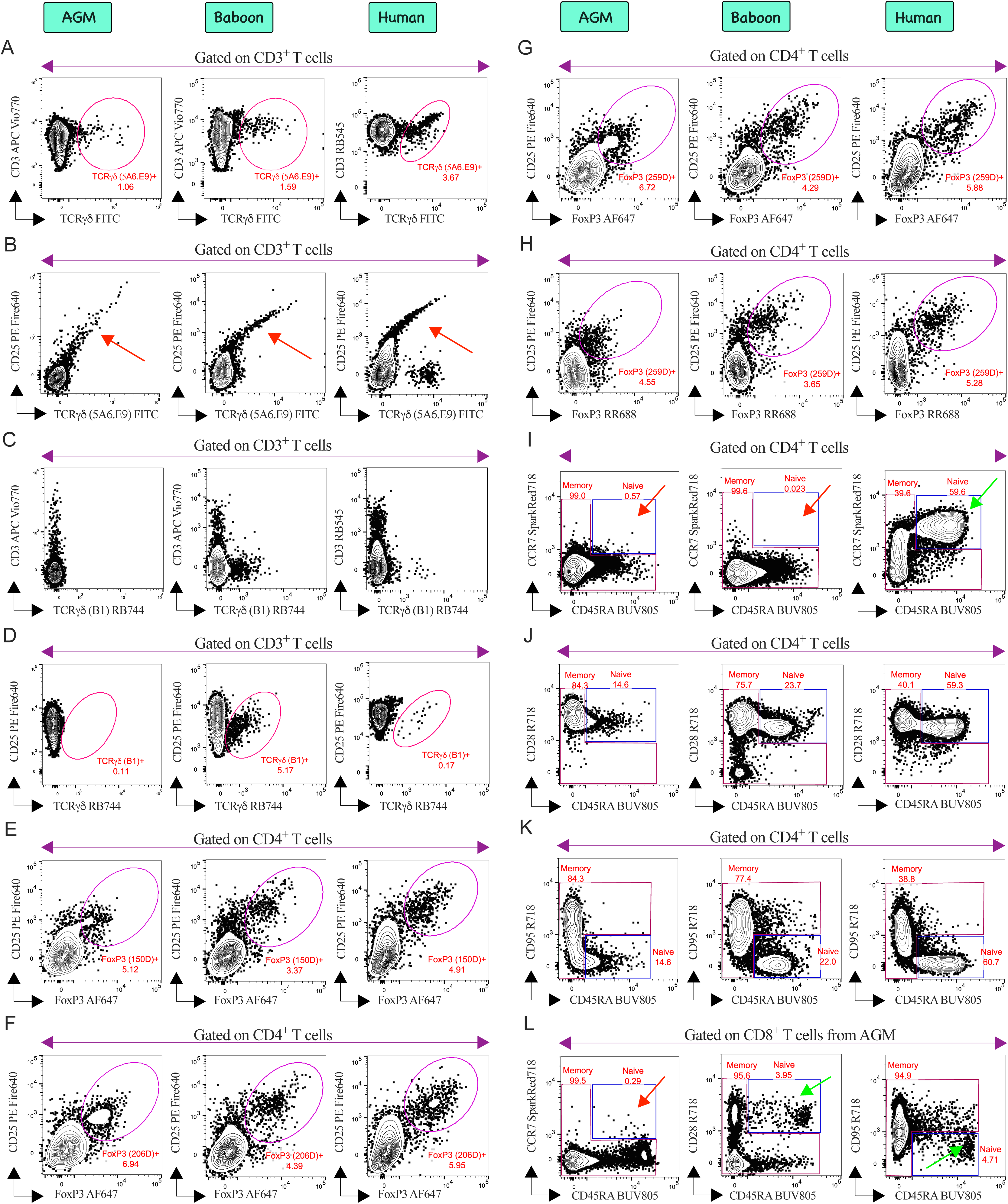
Evaluation of various clones of (A-D) TCRγδ and (E-H) FoxP3 antibodies across NHPs and humans. Based on their performance, specific clones were selected for further analysis. (B) Non-specific binding of TCRγδ clone 5A6.E9 to CD25^+^ T cells was shown in red arrows across all the species, which is absent in (C) TCRγδ clone B1. To assess memory phenotype, three combinations were evaluated: (I) CCR7/CD45RA, (J) CD28/CD45RA, and (K) CD95/CD45RA. The CD95/CD45RA combination was selected for further analysis due to its ability to effectively capture both memory and naive phenotypes. Although (J) CD28/CD45RA successfully captures naive/memory phenotypes across all species, it was not utilized as for antigen-specific memory recall assays, CD28 serves as a co-stimulant. The final panel, L, illustrates the CD8^+^ T cell compartment from AGM PBMCs, validating the memory phenotype strategy. Notably, the AGM T cell compartment is enriched with CD8^+^ T cells rather than canonical CD4^+^ T cells. Red arrows indicated the incompatibility of CCR7 in AGMs to capture naive phenotype. Green arrows indicated the successful capturing of naive CD8^+^ T cell subset. Each column represents a distinct species, as indicated at the top except the final panel L.

**Figure 7.4:**
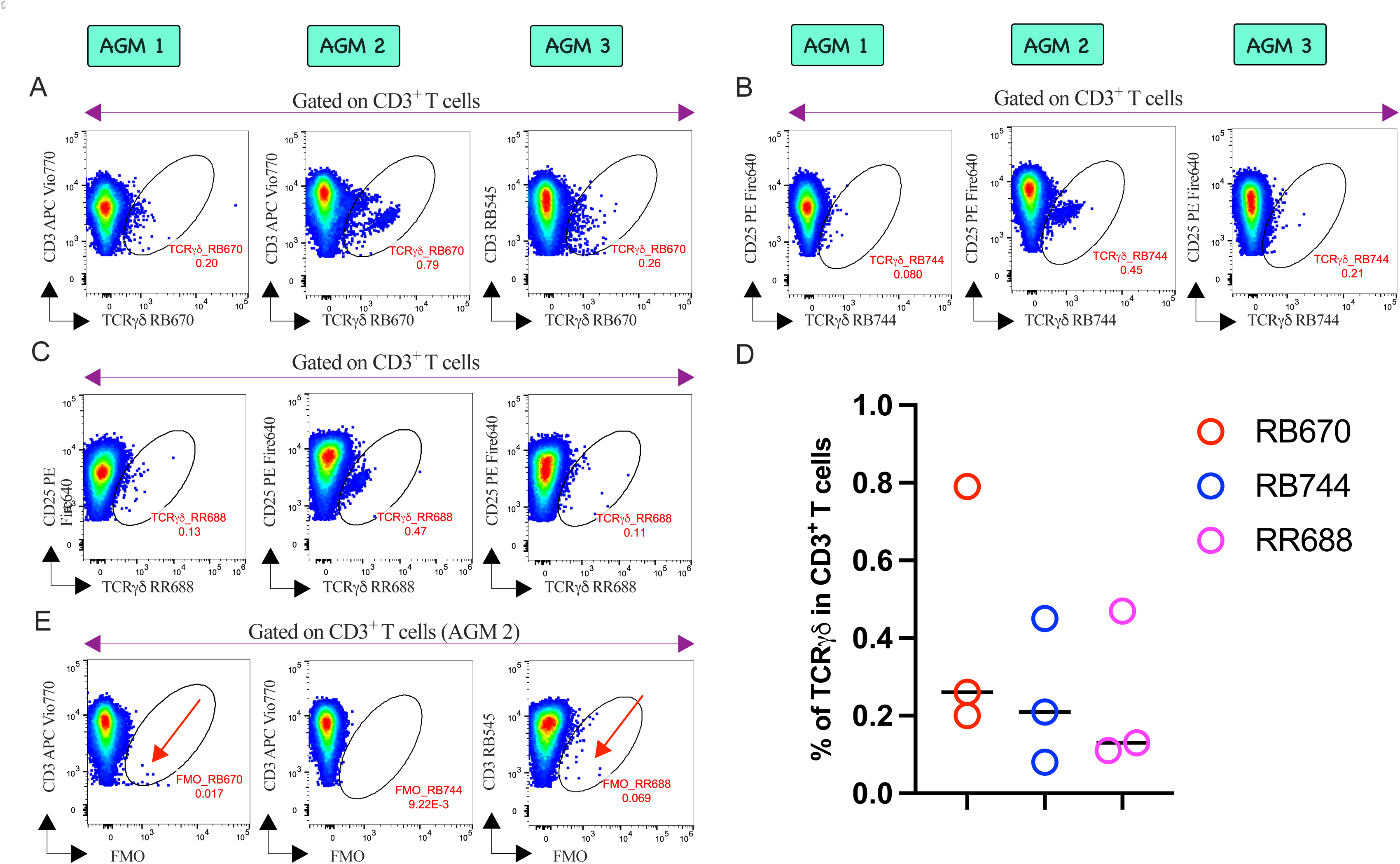
Evaluation of TCRγδ T cells in PBMCs was conducted across three anonymously chosen AGMs. Panel assessments included TCRγδ conjugated to (A) RB670, (B) RB744 and (C) RR688. Although the (D) RB780 conjugate exhibited superior signal intensity, it showed (E) non-specific binding (red arrow) in FMO controls. Conversely, RB744 provided moderate resolution while maintaining target specificity, justifying its inclusion in the optimized panel.

To date, no OMIPs have validated FoxP3 in NHPs. NHP cross-reactivity databases (21) recommend clone 150D for AGMs and baboons, alongside clones 206D and 259D for baboons. Across species, clone 206D demonstrated superior cross-reactivity (Figure 7.3F) compared to 150D (Figure 7.3E) and is extensively documented for profiling FoxP3^+^ Tregs in AGMs (23,42,43), cynomolgus macaques (44,45), and rhesus macaques (46,47), leading us to select 206D for NHP panel. Although both 206D (Figures 7.3F) and 259D (Figures 7.3G) performed well in human PBMCs, 206D is reported to be less sensitive than 259D for detecting steady-state human FoxP3^+^ T cells (48). Because exhaustive clone comparison was beyond the scope of this study, we selected clone 259D for human Treg phenotyping, a choice consistent with its validation in human whole blood in OMIP-114 (49). Validation of clone 259D conjugated to RR688 (Figures 7.3H) demonstrated effective profiling of FoxP3^+^ Tregs in steady-state human PBMCs. However, FoxP3 AF647 was selected for the final panel to prevent deviation of the human panel (3) from its NHP counterpart.

While the CCR7/CD45RA combination was established in OMIP-52 (1) to dissect total memory (central, effector and terminal) T cell phenotypes in rhesus macaques, this strategy failed to detect CCR7^+^ CD45RA^+^ naive cells in NHPs (red arrow, Figure 7.3I), despite resolving them in humans (green arrow, Figure 7.3I). The standard CD28/CD45RA combination (34–36,50) cross-reactively identified naïve and memory T cell populations across all tested species (Figure 7.3J). However, utilizing CD28 as an in vitro co-stimulant (3) necessitated an alternative phenotypic gating strategy. Adapting OMIP-75 (2), which pairs CD95 with CD28 to phenotype NHP memory T cells, we bypassed CD28 by combining CD45RA directly with CD95, successfully delineating naive/memory T cell compartments in both NHPs and humans (Figure 7.3K). To re-validate these gating strategies given the use of recycled animals, we evaluated the CCR7/CD45RA, CD28/CD45RA, and proposed CD95/CD45RA combinations side-by-side in anonymously chosen AGMs (Figure 7.3L). The CCR7/CD45RA combination consistently failed to capture CCR7^+^ CD45RA^+^ naive cells (red arrow, Figure 7.3L), risking naive cell contamination within the memory gate. In contrast, both CD28/CD45RA and CD95/CD45RA combinations reliably identified the naive population as CD28^+^ CD45RA^+^ and CD95^-^CD45RA^+^, respectively (green arrows, Figure 7.3L).

In Figure 8, we validated the gating strategy for challenging fluorochromes and rare cell populations using FMO controls. Rested cryopreserved PBMCs from AGMs (Figures 8A-L) and olive baboons (Figures 8M-X) were analyzed under either steady-state or Con-A stimulated conditions.

**Figure 8:**
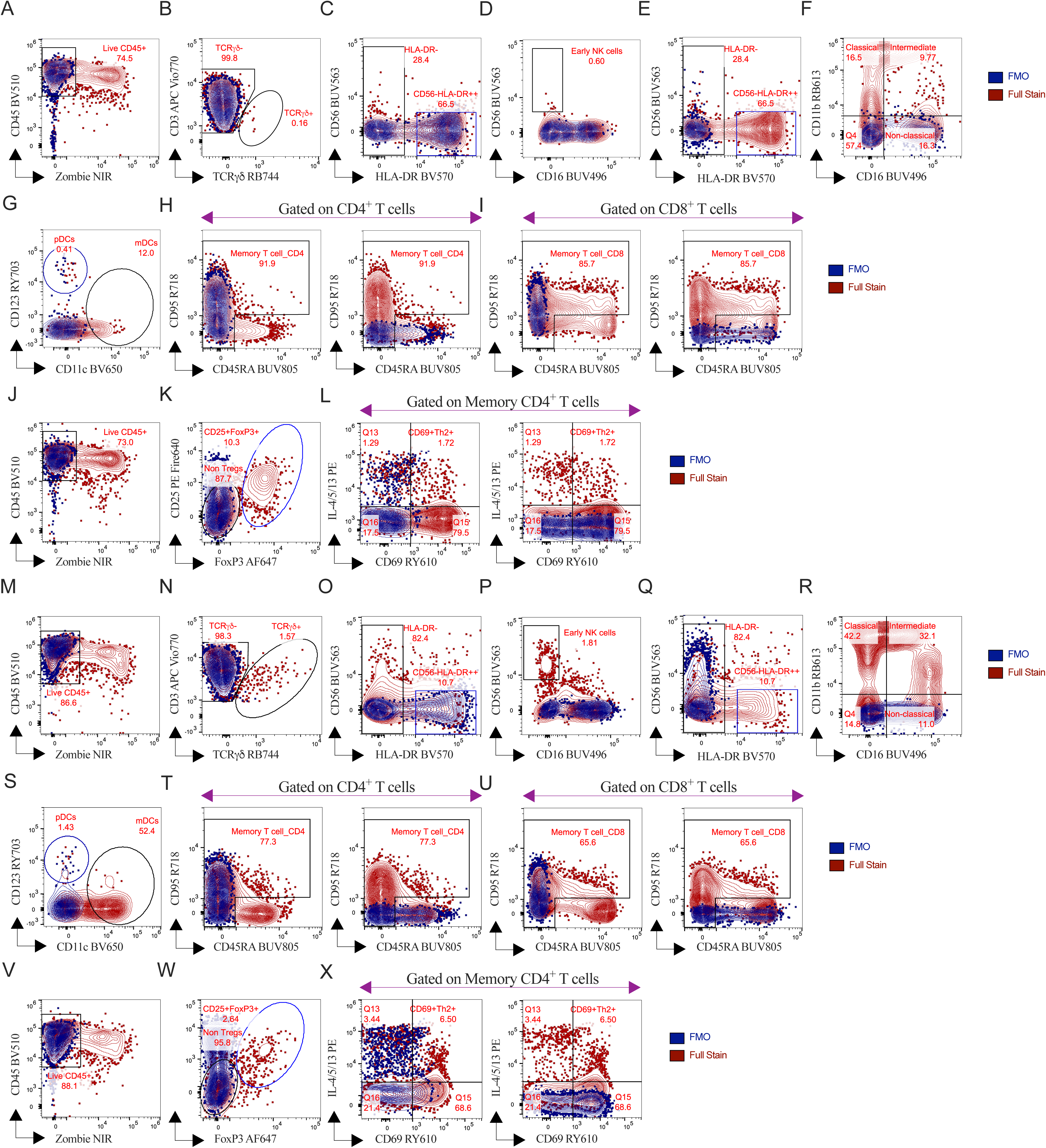
Evaluation of FMO controls. To validate manual gating for challenging fluorochromes and rare populations, this study utilizes FMO controls across (A-L) AGM and (M-X) olive baboon PBMCs. Surface staining for steady-state, unstimulated cryopreserved NHP PBMCs include FMO plots for (A, M) live CD45^+^ cells, (B, N), TCRγδ cells, (C, D & O, P) CD56^+^ subsets, (E, Q) HLA-DR^high^ cells, (F, R) CD11b^+^ cells and (G, S) CD11c^+^ mDCs with memory phenotypes for BUV805 and R718 shown in H, I & T, U. Nuclear staining procedures provide FMO plots for (J, V) live CD45^+^ cells, (K, W) steady-state Treg subsets and (L, X) Con-A stimulated CD69^+^ activated Th2 cells. All FMO controls (blue) overlaid on full stains (brown) for direct comparison.

### 3.3 Single Color Controls, Other Technical Considerations and Data Analysis

#### 3.3.1 Single Color Cellular Controls to Overcome Unmixing Errors

Initially, we evaluated the NHP panel using UltraComp eBeads™ Spectral Unmixing Beads (Table 5) alongside ArC™ Amine Reactive Compensation Beads (Table 6) to calculate the SSM. However, fully stained NHP PBMCs (Figure 9) revealed distinct unmixing artifacts-particularly from Zombie NIR across the violet (Figures 9A, G), blue (Figures 9B, H, C, I), and red spectra (Figures 9D, E, J, K). Mirroring our human panel optimization (3), replacing ArC™ beads with NHP-specific, heat-killed cellular controls effectively resolved these unmixing errors (Figure 9). Similarly, NHP-specific cellular controls were required for HLA-DR BV570 to correct unmixing errors observed with BV421 (Figures 9F, L). To prevent operator bias, we avoided manual overrides via the “unmixing adjuster”, relying solely on the ID7000™ Weighted Least Squares Method (WLSM) algorithm. Finally, treatment-matched NHP unstained controls were incorporated as a dedicated autofluorescence parameter (AF color 1) during unmixing, bypassing the built-in “Autofluorescence Finder” tool given the absence of baseline spectral anomalies.

**Figure 9:**
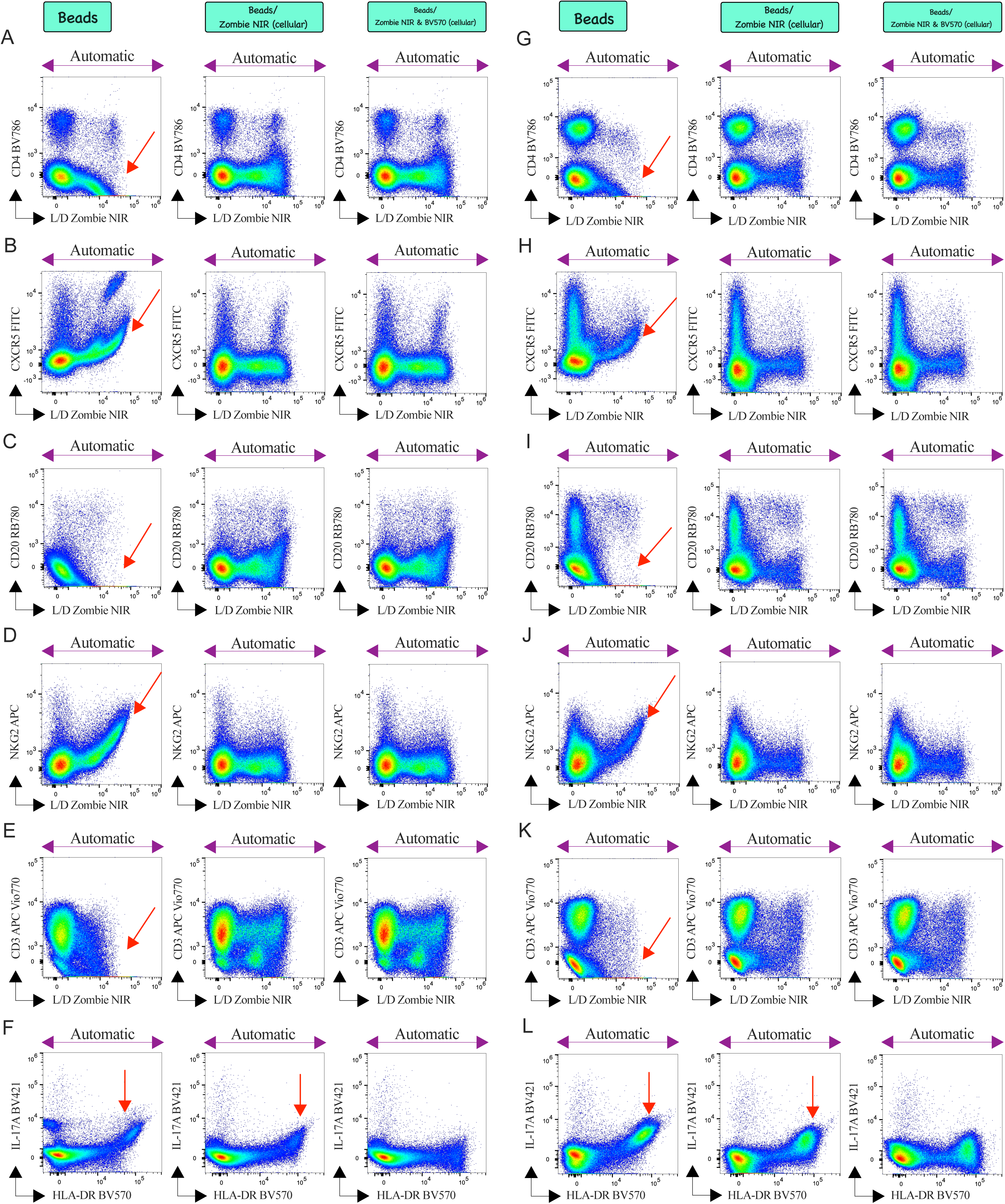
To detect unmixing errors, fully stained Con A-stimulated PBMCs (20 h) from (A-F) AGMs and (G-L) olive baboons were screened using an N x N matrix evaluated on singlets. The process utilized UltraComp Spectral eBeads™ alongside ArC™ reactive beads (indicated on top as “Beads”), with the latter consistently revealing similar unmixing errors (red arrows) across both species. These bead-mediated issues are effectively resolved by substituting ArC™ amine reactive compensation beads with cellular controls (indicated on top as “cellular”), particularly for the Zombie NIR dye. Furthermore, unmixing for (F & L) HLA-DR BV570 using Spectral eBeads highlighted potential errors (red arrows) in NHPs, which were also successfully mitigated by employing cellular controls for that marker. All observations labeled as “automatic” reflect results obtained without manual adjustments via the ID7000™ unmixing adjuster.

##### Step-by-step staining protocol for preparing the reference control for unmixing

The preceding sections adopt the methodology outlined in **Section 3.3.1** of OMIP-XXX-A (3). All acquisitions were performed on an ID7000™ spectral cell analyzer within 16 h of staining using fresh reference controls, collecting 10,000 bead events or 50,000 singlets for Zombie NIR/HLA-DR controls. Spectral unmixing was executed via the WLSM algorithm to compute the SSM (Figure 3).

#### 3.3.2 Other Technical Considerations

##### 3.3.2.1 Stimulant Optimization

Similar to our observations during human panel optimization (3), we initially evaluated two mitogenic stimuli to induce AIM and ICS profiles in NHP PBMCs (Figure 10). Short-term (5 h) stimulation with PMA/ionomycin induced marked CD4 downregulation (red arrows, Figures 10D, I), a phenomenon mirrored in human PBMCs (3), and altered overall CD3^+^ T-cell subset architecture (red arrows, Figures 10B, G). Consequently, PMA/ionomycin was deemed unsuitable for AIM assays (Figures 10E, J), which require extended incubation periods (51). Conversely, 20 h stimulation with Con A maintained CD3^+^ T-cell subset distribution (Figures 10L, Q) without compromising CD4 cell-surface expression (Figures 10N, S). Despite these limitations, PMA/ionomycin was utilized to titrate Th lineage markers (Figures 5.1, 5.2) due to its superior capacity for Th1 induction (Figures 10E, J), whereas Con A was employed for AIM titration (Figures 5.1, 5.2). Ultimately, mirroring the validation strategy of the human panel, we confirmed final panel performance following Con A stimulation in PBMCs from four AGMs and five baboons (comprising three olive baboons and two olive-yellow hybrids) (Figures 11, 12).

A potential limitation of this study is that panel validation was restricted to mitogen stimulation without direct assessment of antigen-specific AIM or ICS. However, because mitogen-driven AIM and ICS responses were robustly detected (Figures 10-12), we reasoned that antigen-specific responses could be similarly resolved. Indeed, in an ongoing clinical trial evaluating an adjuvanted mRNA SARS-CoV-2 vaccine in AGMs, this panel successfully detected SARS-CoV-2 peptide-specific AIM and ICS responses in immunized animals. Furthermore, in an ongoing study examining cellular immunity in *Bordetella pertussis* infected infant baboons, the panel reliably captured AIM and ICS upregulation following stimulation with *B. pertussis* proteins and peptides. These preliminary data from ongoing studies were omitted here as they fall beyond the scope of the present manuscript.

**Figure 10:**
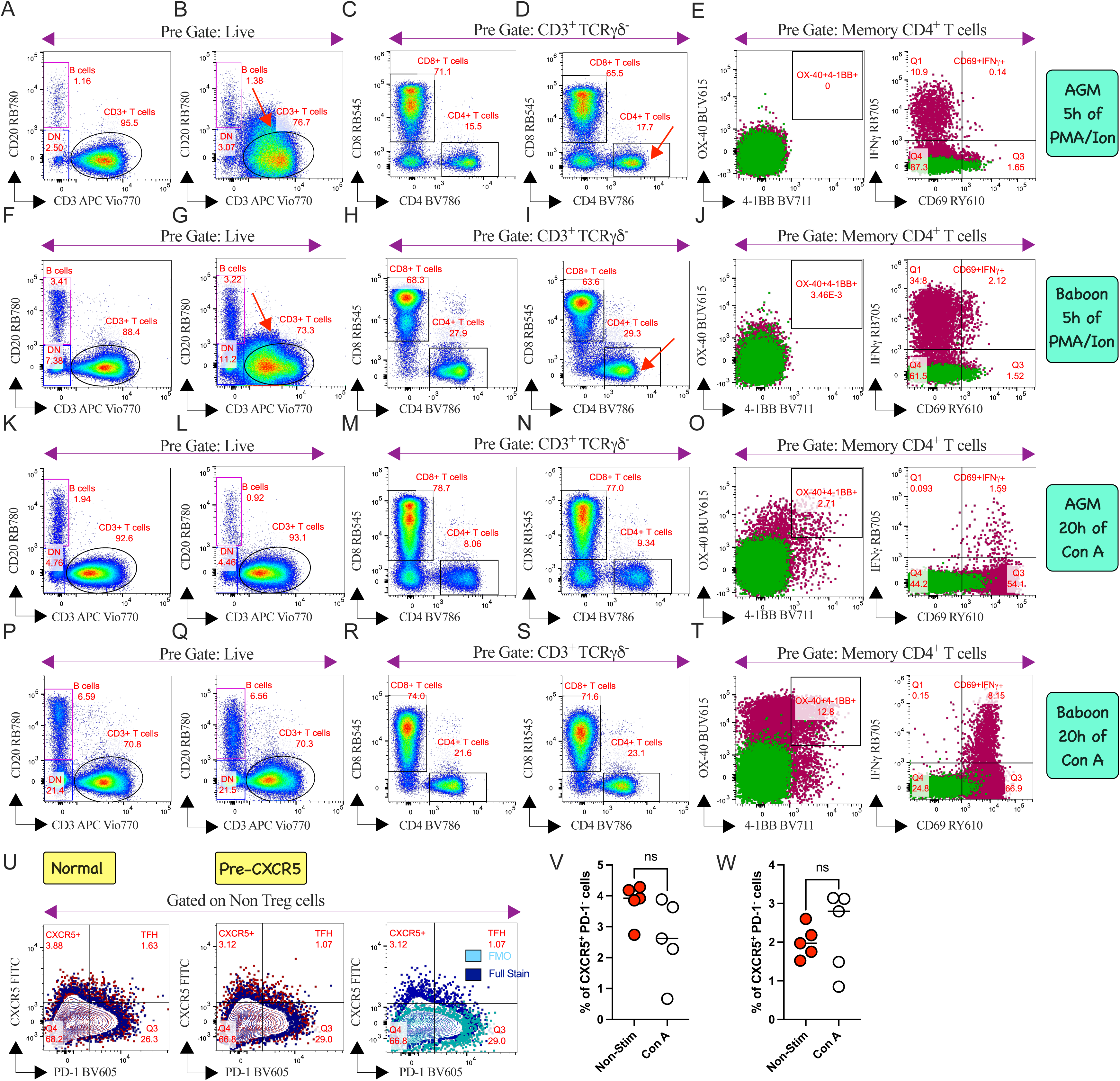
Optimization of stimulant conditions was evaluated using (A-E, K-O) AGM PBMCs and (F-J, P-T) baboon PBMCs, with specific conditions listed at the far right of each row. Bivariate plots illustrate the variance between (A, C, F, H, K, M, P, R) steady-state conditions and their (B, D, G, I, L, N, Q, S) stimulated counterparts across NHPs. Following stimulation with either (A-J) mitogens or (K-T) Con A, (E, J, O, T) AIMs and the Th1 marker IFNγ profile were initially assessed. Short-term mitogen stimulation induced (B, G) CD3^+^ T cell subset distortion and (D, I) CD4^+^ T cell downregulation, as indicated by red arrows. Conversely, (L, N, Q, S) prolonged Con A stimulation produced no such anomalies in NHP PBMCs. Assessment of (U-W) CXCR5 expression dynamics following Con A stimulation was evaluated in baboon PBMCs after nuclear staining, with columns representing distinct immunophenotyping steps as indicated at the top. To evaluate stimulant-dependent CXCR5 alterations, CXCR5 FITC was added either during standard surface staining (“Normal”) or 15 minutes prior to stimulant addition (“Pre-CXCR5”). Activation profiles are visually depicted in (U) representative bivariate plots from an individual animal by overlaying steady-state subsets (blue) onto 20 h Con A stimulated subsets (brown). Additionally, the FMO control subset (cyan) was overlaid on the full stain upon stimulation (blue) to visually validate the gating strategy. Graphical representations of the CXCR5 profile under (U, V) normal or (U, W) pre-CXCR5 conditions after stimulation illustrate the magnitude of stimulant-specific variation within the CD4^+^ T cell compartment. Statistical comparisons were performed using either a two-tailed paired t-test or a Wilcoxon test corrected for multiple comparisons (ns, non-significant; n=5 per group).

**Figure 11:**
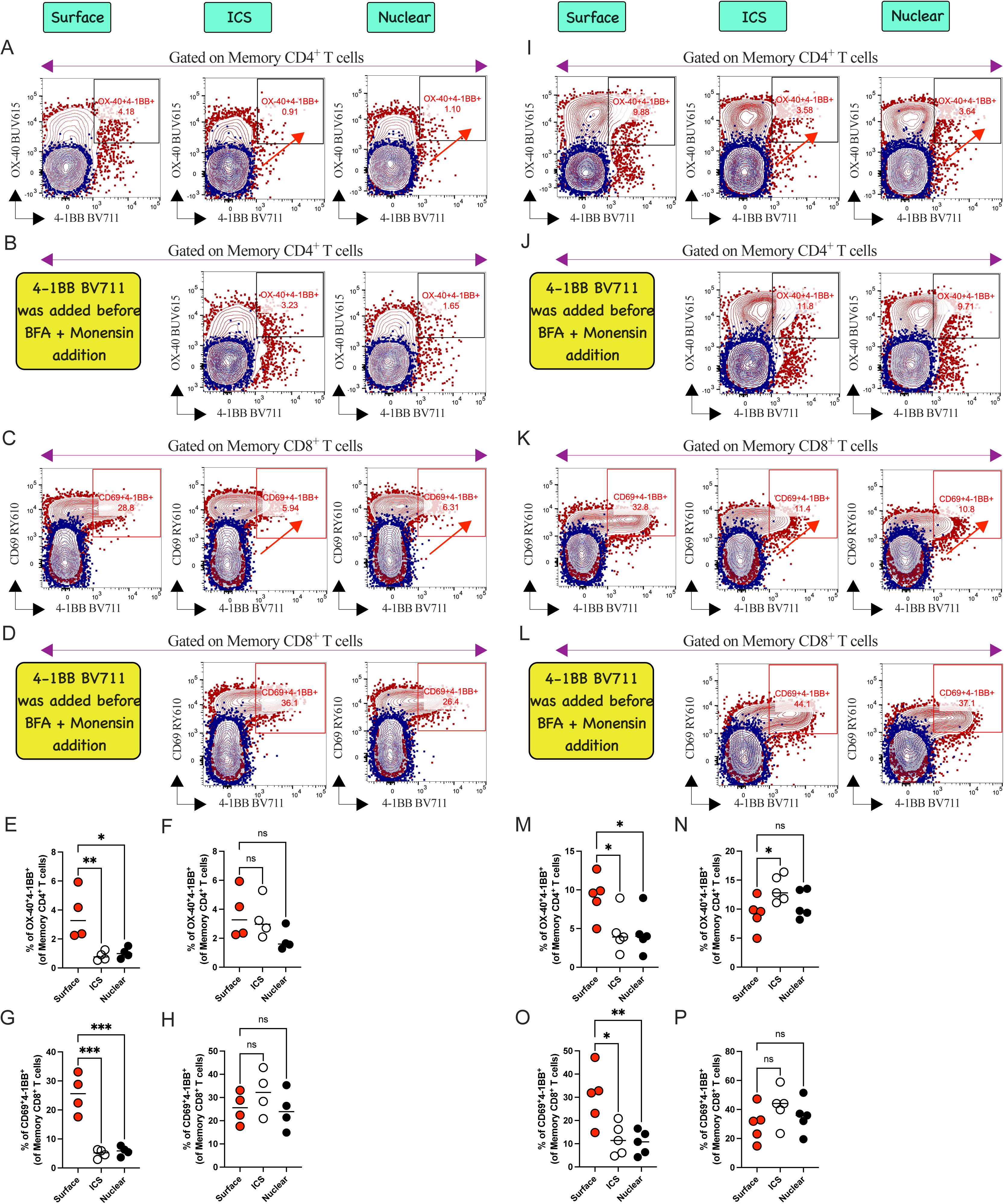
Validation of cytokine blocker mediated downregulation of CD137 (4-1BB) and strategies to overcome it. 4-1BB activation profiles were evaluated in Con A stimulated PBMCs from (A-D) AGMs and (I-L) baboons following surface, ICS and nuclear staining, with columns representing specific immunophenotyping steps as indicated at the top. Activation profiles are visually depicted in representative bivariate plots from an individual NHP by overlaying steady-state subsets (blue) onto 20 h Con A stimulated subsets (brown). Con A induced 4-1BB expression was significantly downregulated during traditional (red arrows; A, C, I, K) ICS and nuclear staining, as quantified in graphical representations for (E, G) AGMs and (M, O) baboons. This downregulation was effectively rescued by introducing 4-1BB BV711 antibody prior to the cytokine-blocking step (indicated by yellow boxes; B, D, J, L), as demonstrated across memory CD4^+^ and CD8^+^ T cell compartments in (F, H) AGMs and (N, P) baboons. Statistical comparisons were performed using either a one-way ANOVA or a non-parametric Kruskal-Wallis test with corrections for multiple comparisons (*P<0.05, **P<0.01, ***P<0.001, ns: non-significant; n=4-5 animals per group, with each circle representing an individual subject).

**Figure 12:**
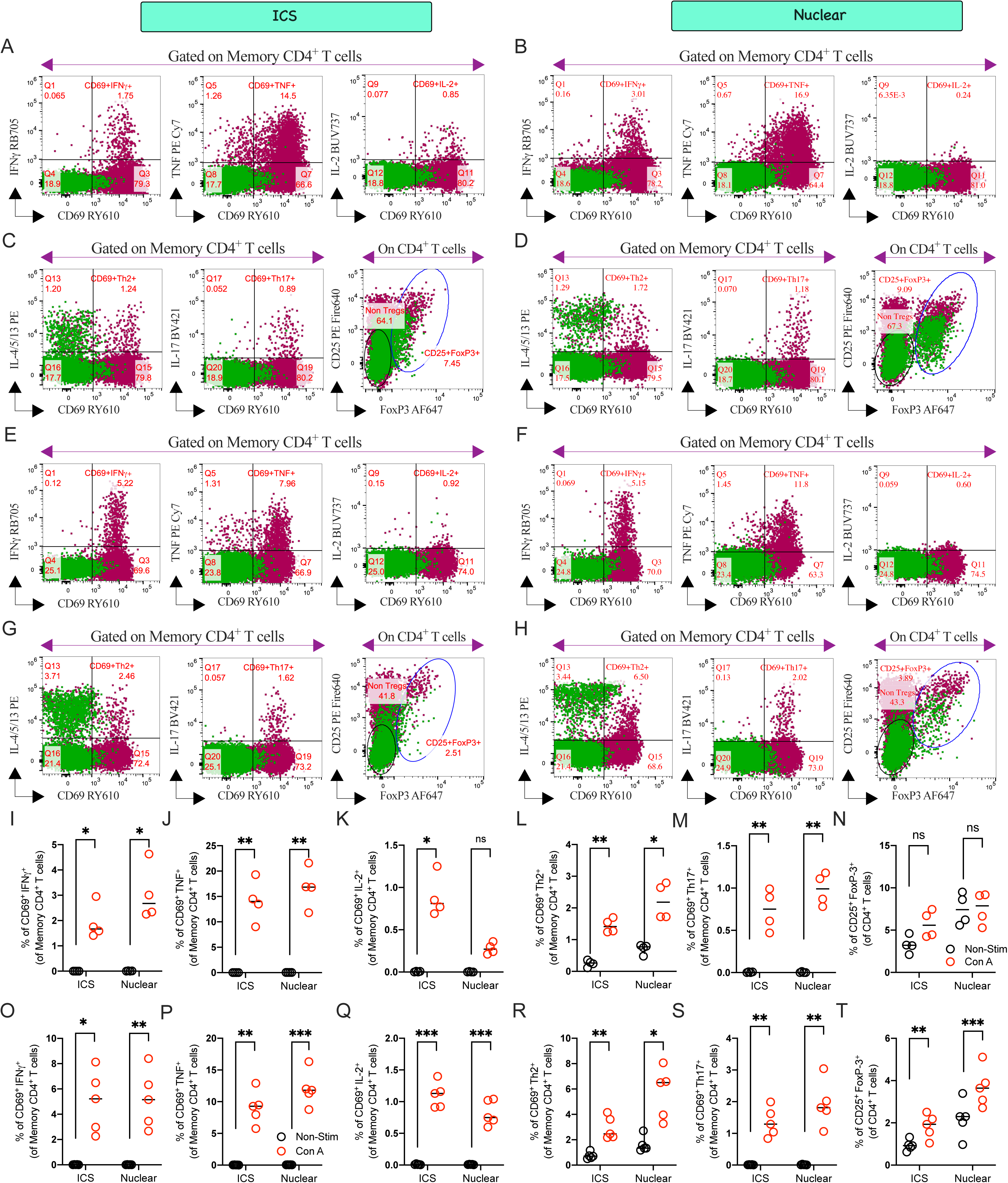
Intracellular cytokine and FoxP3 profiles in Con A stimulated (A-D) AGM and (E-H) baboon PBMCs using ICS or nuclear staining. Th1/Th2/Th17 and FoxP3 signatures were evaluated following (A, C, E, G) ICS or (B, D, F, H; indicated at top) nuclear staining. Representative bivariate plots from individual animals depict activation dynamics by overlaying steady-state subsets (clover) onto 20 h Con A stimulated subsets (cayenne). Quantification of Th1/Th2/Th17 and Treg profiles in Con A stimulated (I-N) AGM and (O-T) baboon PBMCs using ICS or nuclear staining. Statistical comparisons were performed using either a two-tailed paired *t*-test or Wilcoxon test corrected for multiple comparisons (*P<0.05, **P<0.01, ***P<0.001, ns: non-significant; n=4-5 animals per group, each circle represents an individual animal).

##### 3.3.2.2 Sensitivity of CXCR5 and AIMs during ICS and Nuclear Stain

Stimulus-induced CXCR5 downregulation was previously reported in rhesus macaques (1), and we observed the same effect in AGMs following Con A stimulation, though not in humans (3). Con A similarly downregulated CXCR5 in baboons (Figures 10U, V). To address this, we implemented the “Pre-CXCR5” strategy used in our human panel (3), adding the CXCR5 FITC antibody 15 minutes prior to stimulation and incubating at 37°C. This pre-incubation step successfully prevented loss of surface expression and allowed us to capture Con A-driven CXCR5 upregulation in baboons (Figures 10U, W). We also observed cytokine blocker-specific (brefeldin A and monensin) downregulation of 4-1BB during both ICS and nuclear staining in AGMs (Figures 11A, C, E, G) and in baboons (Figures 11I, K, M, O). To overcome this, we applied the strategy validated in our human panel (3), successfully rescuing surface expression by staining with anti-4-1BB BV711 prior to adding cytokine blockers (Figures 11B, D, F, H, J, L, N, P). Notably, no protocol-specific differences (ICS vs. nuclear staining) in the AIM profile were observed (Figures 11E-H, M-P).

##### 3.3.2.3 Differences in Cytokine Profile between ICS and Nuclear Stain

Following our human panel design (3), we evaluated how protocol-specific workflows-specifically ICS versus nuclear staining-impacted the resolution of rare Th1, Th2, and Th17 subsets post-Con A stimulation (Figure 12). Such direct, head-to-head protocol comparisons using NHP PBMCs are rarely documented. Both staining strategies effectively captured Th1 profiles within memory CD4^+^ T cells following Con-A stimulation, with the exception of lower IL-2 detection during nuclear staining in AGMs (Figure 12K). We did not further investigate the reduced IL-2 detection observed in AGMs following nuclear staining for several reasons: first, these were re-used (recycled) animals; second, this marked loss of IL-2 was not observed in human (3) or baboon samples (Figure 12Q); and third, resolving Th1 marker IFNγ and Treg signatures remained our primary objective, consistent with our human panel design (3). Nonetheless, researchers intending to measure IL-2 in AGMs using a nuclear staining protocol should exercise caution. Nuclear staining demonstrated superior efficiency in capturing IFNγ (Figure 12O) and TNF (Figure 12P) signatures in baboons. Both ICS and nuclear staining demonstrated comparable efficacy in capturing Con A-stimulated Th2 (Figure 12L, R) and Th17 profiles (Figure 12M, S) across NHPs. As anticipated, nuclear staining exhibited optimal efficiency in targeting FoxP3^+^ Tregs (Figures 10D, H, N, T) within the CD4^+^ T cell compartment across both species, whereas ICS resulted in suboptimal Treg capture (Figures 10C, G, N, T).

In summary, we established and optimized a comprehensive immunophenotyping panel capable of simultaneously mapping AIMs, Th1/Th2/Th17 lineages, TFH subsets, and Tregs across AGMs, baboons, and humans. We envision that this versatile spectral tool will serve as a robust framework for translational researchers aiming to interrogate previously uncharted cellular landscapes in NHP models.

#### 3.3.3 Data Analysis

Experimental procedures for sample acquisition (Sony ID7000™), spectral unmixing, gating strategy (FlowJo v10.10.1), and high-dimensional visualization (OMIQ) strictly mirror those described in OMIP-XXX-A (3) and are not repeated here.

## 4 Data Availability Statement

All data supporting the findings of this study are available within the article. NHP-specific unmixed, quality-controlled FCS files and FlowJo workspace (WSP) files are deposited in Zenodo (https://doi.org/10.5281/zenodo.23672066). The deposited data will be made publicly available upon publication. Repository metadata mapping individual FCS filenames to their corresponding stimulation conditions are provided in a “Sample Description” PDF generated from workspace screenshots. Panel optimization datasets (antibody titrations, FMOs, clone validation, ICS profile, related to Figures 7-12) are available from the corresponding authors upon reasonable request.

## Author Contributions

**Soumik Barman:** conceptualization, methodology, data curation, investigation, validation, formal analysis, visualization, writing-original draft, writing-review and editing.

**Arsh Patel:** conceptualization, software, validation, formal analysis, visualization, data curation, writing-review and editing.

**Michael J. Buonopane:** software, validation, formal analysis, visualization, data curation, writing-review and editing.

**Ben Janoschek:** conceptualization, software, validation, formal analysis, visualization, writing-review and editing.

**Ravi Hingorani:** conceptualization, validation, writing-review and editing.

**David J. Dowling:** conceptualization, supervision, funding acquisition, project administration, resources, writing-review and editing.

## Acknowledgements

Partial support for the Precision Vaccines Program (PVP) is provided by Boston Children’s Hospital, Department of Pediatrics. For their valuable discussion and support, the authors wish to thank the members of the PVP, Dr. Etsuro Nanishi (former member, PVP), Dr. Ofer Levy (Program Director, PVP), Suzan Lazo (Director, Cytometry Cores, DFCI), and Matthew Powers (Assistant Director, Cytometry Cores, DFCI). Appreciation is also extended to the late Dr. Andrew Draghi II (Sony Biotechnology Inc.) for his valuable insights into the Sony ID7000™ Spectral Cell Analyzer, and to Dr. Rajesh Panigrahi (Waters Biosciences) for his expert input on the Real Blue/Yellow fluorochromes during panel optimization. Dr. Hanne Andersen (Bioqual Inc.) and Deborah Chavez (SNPRC) are also thanked by Soumik Barman for their technical assistance and helpful feedback. The authors also thank Dr. John Quinn (Waters Biosciences) for his technical guidance on the FlowJo data analysis. Lastly, the authors gratefully acknowledge Eric Haas (Ionic Cytometry Solutions) for conducting quality assurance on the deposited datasets.

## Supporting Information

Additional supporting information can be found online in the Supporting Information section. **Data S1:**

Cytometry Part A Author Checklist: MIFlowCyt-Compliant Items.

